# A light-labile signal molecule acts as a photoregulator of secondary metabolite biosynthesis in a heterotrophic bacterium

**DOI:** 10.64898/2026.03.02.708258

**Authors:** Alissa Dierberger, Magdalena Rose, Raimund Nagel, Torsten Jakob, Charles Simpson, Shuaibing Zhang, Vivien Hotter, Maria Mittag, Pierre Stallforth, Qing Yan, Severin Sasso

## Abstract

While light is essential for many phototrophic organisms, our knowledge of the light-dependent regulation in heterotrophic organisms is scarce. Here, we found that the heterotrophic soil bacterium *Pseudomonas protegens* differentially accumulated 234 secreted metabolites depending on the light conditions. These metabolites included important antimicrobials such as pyoluteorin, 2,4-diacetylphloroglucinol, pyrrolnitrin, and rhizoxin analogs. Pyoluteorin, for instance, was 140-fold more abundant in the dark, due to a strong upregulation of the transcript of the pyoluteorin biosynthesis gene *pltL*. We discovered that 2,4-dichlorophloroglucinol (PG-Cl_2_), an activator of pyoluteorin biosynthesis, acts as a photoregulator that degrades much faster in the presence of light than in the dark. The resulting higher PG-Cl_2_ levels activate the *pltL* promoter in the dark. These findings reveal a novel mode of regulation in which light instability of a signal molecule allows the producing organism to sense and transmit light information to regulate the biosynthesis of a secondary metabolite.

## Introduction

Light shapes our environment as a source of energy and information for phototrophic organisms, but also influences non-phototrophic organisms^1^. In recent years, it has been shown that light can have major effects on the lifestyles of non-phototrophic organisms by regulating various processes such as metabolic pathways, growth, motility or virulence^2^. Photoreceptors along with their chromophores were found to modulate the behavior of several soil-dwelling and plant-associated heterotrophic bacteria^2^. For example, *Pseudomonas syringae* pv. *tomato* DC3000 has one protein with a light-oxygen-voltage (LOV) domain and two bacteriophytochromes that influence swarming motility and virulence^3^. This strain belongs to a large and diverse genus of Gram-negative bacteria that consists of more than 300 species^4^. *Pseudomonas* spp. are ubiquitous in water, soil and on plants, and their interactions with other organisms range from beneficial to pathogenic^5^. For instance, *Pseudomonas protegens* Pf-5, a strain formerly classified as *Pseudomonas fluorescens*^6^, was isolated from the rhizosphere of cotton seedlings and is used as a biocontrol strain for plant protection^7,8^. This strain produces a variety of secondary metabolites that antagonize phytopathogenic fungi, oomycetes, bacteria, and insects^7,9-12^. In addition, some of these secondary metabolites harm the unicellular soil alga *Chlamydomonas reinhardtii* by inhibiting its growth, causing morphological changes and an increase in cytosolic Ca^2+^, resulting in loss of motility^13-15^.

An important secondary metabolite of *P. protegens* against phytopathogenic microorganisms is pyoluteorin, a hybrid polyketide/nonribosomal peptide. A central part of the pyoluteorin biosynthetic gene cluster is the *pltLABCDEFG* operon that encodes the biosynthetic enzymes^16^ (Fig. 1). Pyoluteorin biosynthesis starts with the activation of L-proline by the L-prolyl-AMP ligase PltF followed by the attachment to the peptidyl carrier protein PltL^17^. After the pyrrolidine ring is oxidized by the acyl-coenzyme A dehydrogenase PltE^17^, the resulting pyrrole is dichlorinated by a single FADH_2_-dependent halogenase, PltA^18^. The type I polyketide synthases PltB and PltC, the thioesterase PltG, and the aromatizing dehydratase PltD participate in the remaining biosynthetic steps, including the formation of pyoluteorin’s resorcinol moiety^19-22^ (Fig. 1). The substrate of the predicted transporter PltIJKNOP has not been identified, and it has been speculated that this transporter functions as an exporter that decreases the intracellular levels of a metabolite that represses pyoluteorin biosynthesis^23^.

**Fig. 1.**
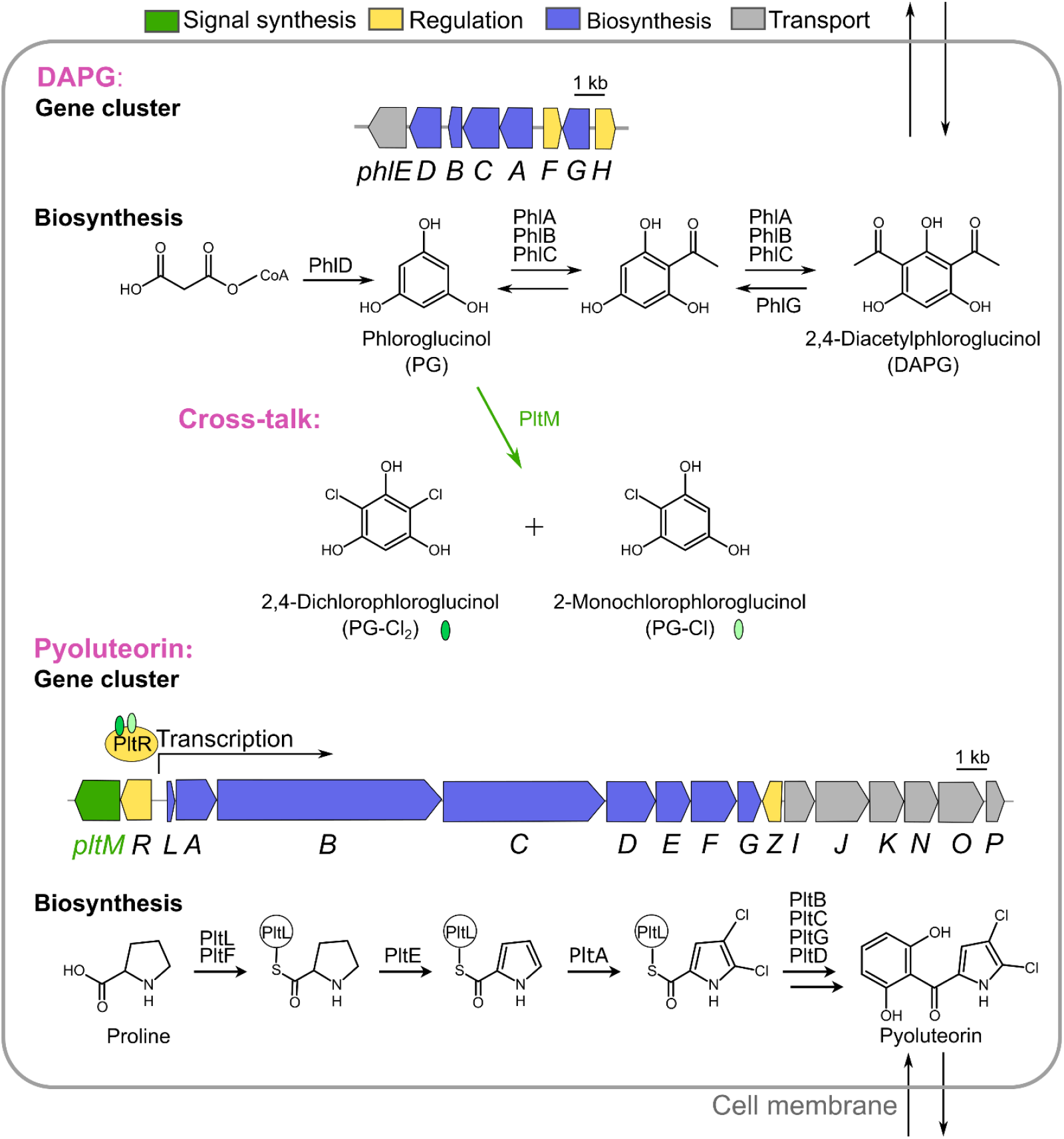
Pyoluteorin biosynthesis and its co-regulation with DAPG biosynthesis. The *pltLABCDEFG* operon in the pyoluteorin (*plt*) gene cluster encodes the biosynthetic enzymes required for pyoluteorin biosynthesis. The halogenase PltM chlorinates PG, a biosynthetic precursor of DAPG. The signal molecules PG-Cl and PG-Cl_2_ induce transcription from the *pltL* promoter, probably by interacting with the transcriptional activator PltR. See text for additional details. (adopted from ref. 16)

The GacS/GacA two-component signal transduction system, consisting of the GacS sensory kinase and the GacA response regulator, acts as a global regulator for the biosynthesis of pyoluteorin and various other secondary metabolites and secreted enzymes^24,25^. Spontaneous mutations in *gacS* or *gacA* may lead to a drop in the production of pyoluteorin, proteases, and other extracellular substances^24,26^. Furthermore, pyoluteorin biosynthesis is regulated in conjunction with the biosynthesis of the secondary metabolite 2,4-diacetylphloroglucinol (DAPG). A biosynthetic precursor of DAPG, phloroglucinol (PG), is chlorinated by the halogenase PltM encoded by the pyoluteorin biosynthetic gene cluster^16^ (Fig. 1). The chlorinated PGs, 2-monochlorophloroglucinol (PG-Cl) and 2,4-dichlorophloroglucinol (PG-Cl_2_), were proposed to interact with the transcriptional activator PltR to activate the *pltL* promoter^16^. While PltR is essential for pyoluteorin biosynthesis, a second transcription factor, PltZ, is additionally required for autoinduction together with PltR (ref. 23). Other factors such as the global regulator Vfr (ref. 27) also regulate pyoluteorin accumulation.

Although it is known that light can influence the lives of heterotrophic bacteria, our knowledge of its influence on secondary metabolism is limited. Here, we find that light and darkness differentially affect the accumulation of many secreted metabolites in *P. protegens*. In view of the numerous studies on the biosynthesis and regulation of pyoluteorin, our finding of pyoluteorin accumulation in the dark but not in light is surprising, and we elucidate a regulatory mechanism that relies on the light instability of an activation signal.

## Results

### Light influences the accumulation of secreted metabolites

In previous work, we cultured *P. protegens* and the photosynthetic microalga *C. rienhardtii* in algal Tris-acetate-phosphate (TAP) medium to characterize the antagonistic effects of *P. protegens* against the alga^13-15^. In this colorless medium, we observed that axenic *P. protegens* cultures turn brownish in continuous darkness whereas the cultures turn white when grown in continuous light (Supplementary Fig. 1a). Since centrifugation of the cultures indicated that the brownish color is caused by secreted metabolites, we extracted these metabolites from the spent medium using ethyl acetate. Similar to the colors of the cultures, concentrated extracts from the spent medium of cultures grown in the light were light brown while concentrated extracts from cultures grown in the dark were dark brown (Supplementary Fig. 1b).

Liquid chromatography-mass spectrometry (LC-MS) was used to elucidate the impact of light and darkness on the exometabolome of *P. protegens*. For this purpose, cultures inoculated with dark-grown pre-cultures were grown in light or darkness for 3 days, and the exometabolomes were subjected to untargeted analysis in one-day intervals. Bacterial growth was not influenced by the light conditions (Supplementary Fig. 1c). A principal component analysis (PCA) shows that the exometabolomes differed very little after 24 h in light or darkness, whereas substantial differences emerged after 48 h (Fig. 2a). Across all timepoints, 735 secreted metabolites were identified, with the reliability of the identification being classified from level 1 (high) to level 5 (low) (see Methods for details). Of these 735 metabolites, 234 (32%) were at least twofold more abundant in light or darkness at one or more timepoints (Fig. 2b, Supplementary Fig. 1d-f and Supplementary Table 1). For example, 37 secreted metabolites were more abundant in light than in darkness while 34 metabolites were exclusively detected in the light after 48 h (Fig. 2b). Conversely, 54 metabolites had an increased concentration in darkness and 50 metabolites were only found in darkness, while the concentrations of 272 metabolites did not differ between light and darkness (Fig. 2b).

**Fig. 2.**
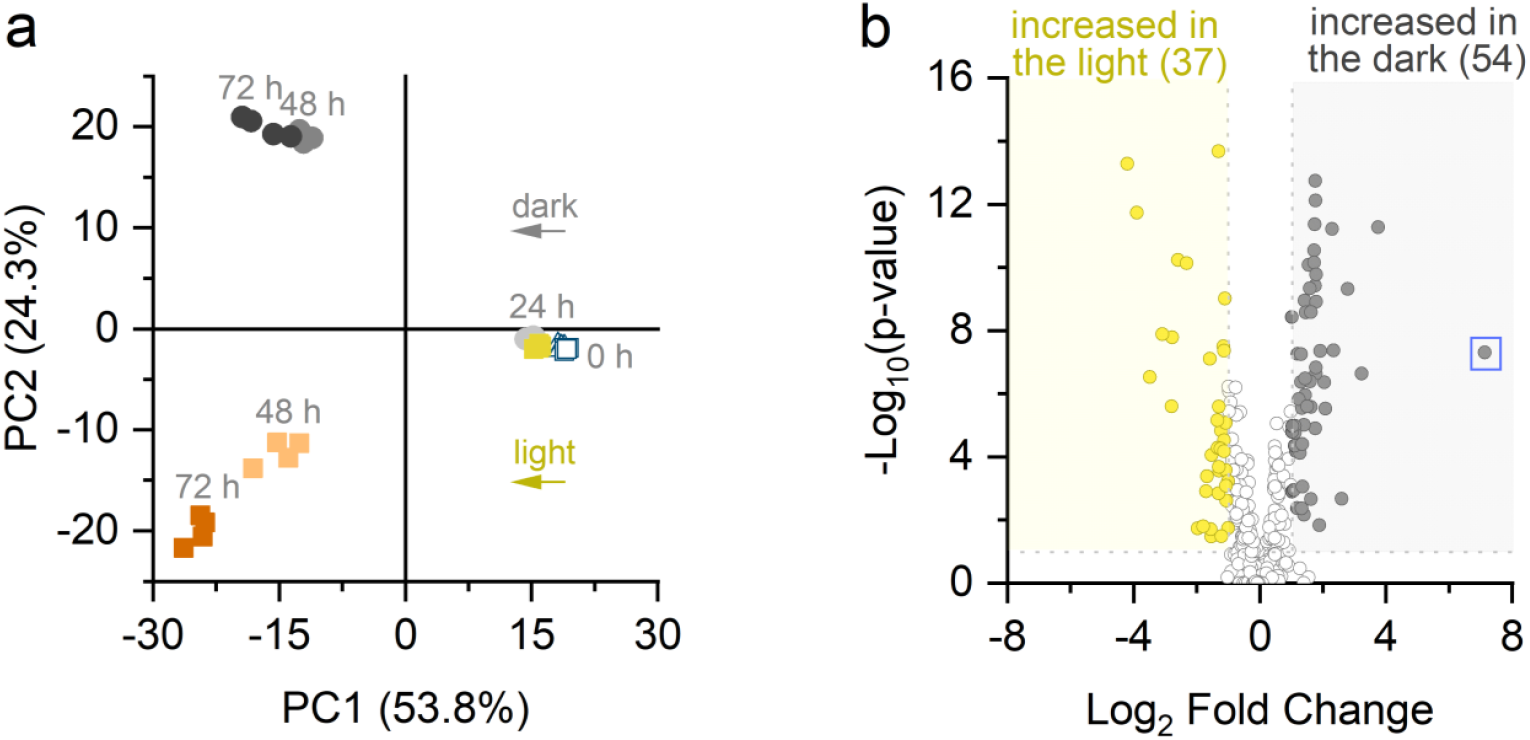
Influence of light and darkness on the *P. protegens* exometabolome. The spent medium of cultures grown in white light of 40 µmol photons m^-2^ s^-1^ or darkness was analyzed 0, 24, 48, and 72 h after inoculation by LC-MS (4 biological replicates each). **a**, Principal component analysis. Principal component 1 (PC1) primarily visualizes differences in metabolite composition caused by time while PC2 primarily visualizes differences caused by light or darkness. **b**, Volcano plot depicting changes between light and darkness after 48 h. The change in metabolite level was considered significant if |log_2_ fold change| ≥ 1 and *p* < 0.05. In addition to the 63 metabolites that were present in higher concentrations in the light than in the dark (shown in the volcano plot), 34 metabolites were only detected in the light and are not shown in the volcano plot. Conversely, 35 metabolites were more abundant in the dark (shown in the volcano plot) while 50 metabolites were only detected in the dark (not shown). Pyoluteorin is highlighted with a blue square. (see Supplementary Fig. 1d,e for volcano plots after 24 h and 72 h and Supplementary Table 1 for a list of differentially accumulated metabolites)

The bacteria accumulated the metabolites rhizoxin S2 (#1), pyoluteorin (#2), 2,4-diacetylphloroglucinol (DAPG, #66), and pyrrolnitrin (#230) in higher amounts in darkness compared to light at one or more timepoints (Supplementary Table 1). The correct identification of these four metabolites was confirmed with standards. Other secreted metabolites accumulated in the dark include the predicted enantio-pyochelin (#15), a siderophore^28^, monodechloroaminopyrrolnitrin (#5), a biosynthetic precursor of pyrrolnitrin^29^, orfamide A (#42), and several indole derivatives (#69, #75, #225). Two putative quinolone derivatives, which are potentially involved in quorum sensing^30^, were also accumulated in darkness (#180) or light (#148).

Among the differentially accumulated metabolites, rhizoxin S2 showed the largest number of derivatives and the highest abundance based on its maximal mass-spectrometric peak area (Supplementary Table 1). Molecular networking based on shared MS^2^ fragments revealed a total of 32 rhizoxin derivatives (Supplementary Fig. 2). This wide variety of rhizoxins can be attributed to their sensitivity to light and oxygen^31,32^ as well as the capacity of *P. protegens* to produce various rhizoxins^33,34^. The differentially accumulated metabolite with the second highest maximal peak area, pyoluteorin, was 140-fold more abundant in darkness after 48 h (blue square in Fig. 2b). Given the well-studied biosynthesis of pyoluteorin^16-23^, the dark-induced accumulation of pyoluteorin was surprising and prompted us to explore this result in more detail.

### Blue light regulates pyoluteorin accumulation

To analyze the possible degradation of pyoluteorin by light, the *in vitro* stability of the metabolite was quantified via LC-MS after an incubation in continuous white light or darkness in TAP medium. Over a period of 48 h, about 30% of pyoluteorin degraded, and while this degradation was slightly faster in the light compared to darkness (Fig. 3a), this difference is too small to explain the 140-fold difference observed by exometabolomics.

**Fig. 3.**
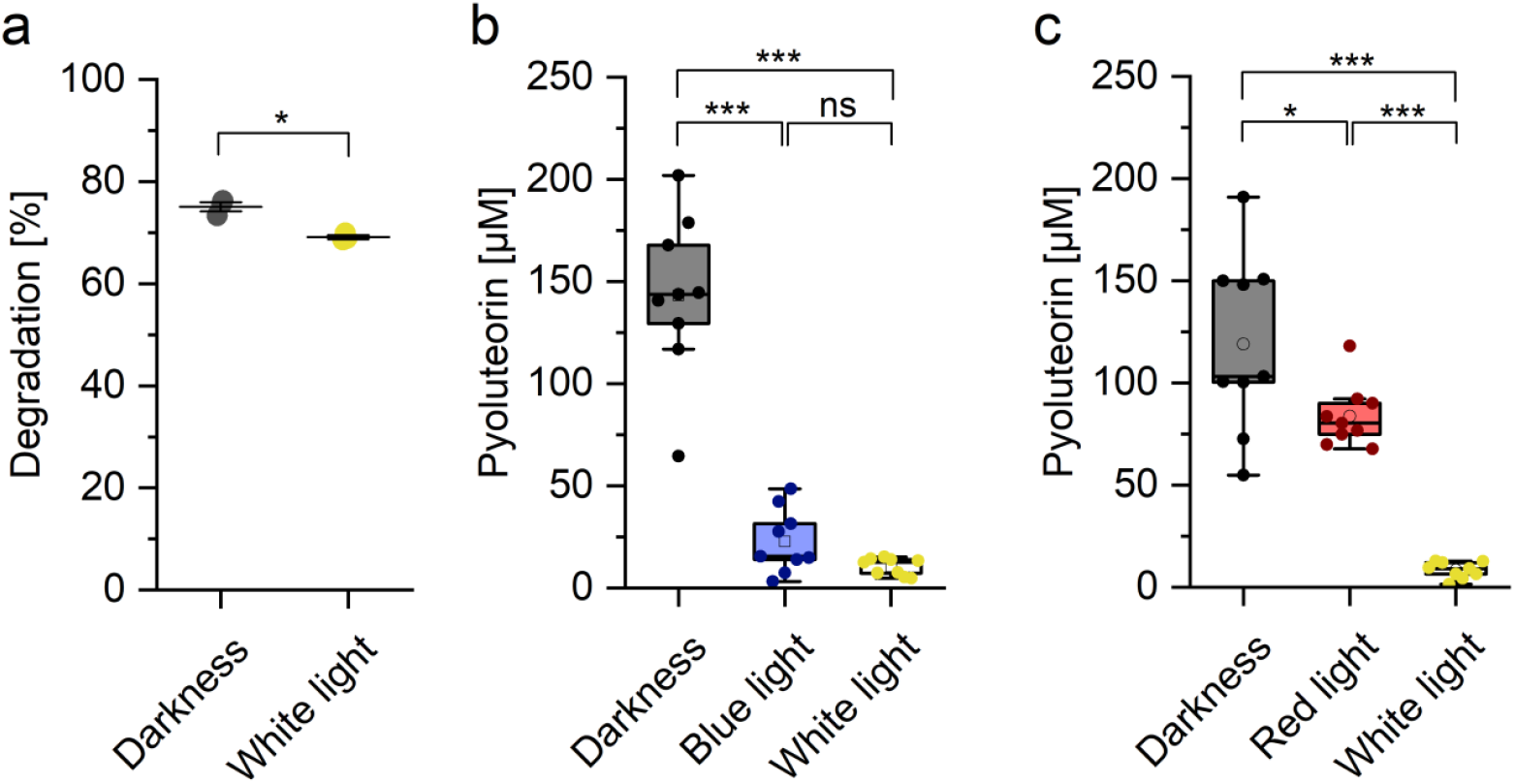
Darkness induces pyoluteorin accumulation in the spent medium of *P. protegens*. **a**, *In vitro* stability of pyoluteorin in darkness and light. 75 µM pyoluteorin in TAP medium were incubated at 20 °C in darkness or white light of 40 µmol photons m^-2^ s^-1^. Recovery (%) is the ratio of the LC-MS signals after 48 h and 0 h. The central horizontal line denotes the mean value and the whiskers denote the standard deviation. An unpaired two-tailed Student’s t-test was used for statistical analysis (*, *p* < 0.05; 3 technical replicates). **b, c** Pyoluteorin accumulation in the spent medium of *P. protegens* grown for 48 h in darkness, blue light, red light or white light (two independent datasets). The light intensities were 40 µmol photons m^-2^ s^-1^ (white light) or 6 µmol photons m^-2^ s^-1^ (blue light and red light). The spent medium of the cultures was extracted with ethyl acetate and pyoluteorin quantified via HPLC. In both **b** and **c**, the experiments were performed three times independently, with three biological replicates each. Filled circles represent data points and open squares or circles represent mean values. The boxes of the box plots denote the interquartile range, the central horizontal line denotes the median, and the whiskers denote the minimal and maximal values. A one-way ANOVA with a Tukey multiple-comparison *post-hoc* test was used for statistical analysis (*, *p* < 0.05; ***, *p* < 0.001; ns, not significant).

To examine whether different light qualities affect the accumulation of pyoluteorin, *P. protegens* was cultivated under blue, red or white light and the concentration of pyoluteorin was determined in the spent medium by high-performance liquid chromatography (HPLC). In two independent datasets, growth under white light for 48 h resulted in a 14-fold lower pyoluteorin concentration in the spent medium compared to growth under darkness (Fig. 3b,c), qualitatively confirming the results from exometabolomics. Similar to white light, blue light strongly inhibited the accumulation of pyoluteorin (Fig. 3b). In contrast, red light had a minor influence and only decreased the concentration of pyoluteorin in the spent medium by 1.4-fold compared to cultures grown in darkness (Fig. 3c). In both datasets, the cell densities of the bacterial cultures were similar and showed no statistically significant differences after 48 h, excluding any relevant effects of the cell densities on pyoluteorin concentrations (Supplementary Table 2). Taken together, these results indicate that pyoluteorin is light-stable over several days, indicating that the differential accumulation of this metabolite under blue light, red light or in darkness is regulated by the bacteria.

### Transcription of *pltL* is upregulated in darkness

*P. protegens* and other *Pseudomonas* spp. are known to accumulate spontaneous mutations in the GacS/GacA regulatory system, and mutations in *gacS* or *gacA* can decrease the production of pyoluteorin, proteases, and other extracellular substances^24,26^. Cultivation on skim milk agar (SMA) plates indicated that *P. protegens* secretes proteases both in light and darkness (Supplementary Fig. 3), suggesting that the GacS/GacA system is functional under both conditions and that the dark-induced accumulation of pyoluteorin is regulated by another mechanism. To test whether the accumulation of extracellular pyoluteorin is regulated at the level of biosynthesis, we used reverse transcription-quantitative real-time PCR (RT-qPCR) to quantify the transcript levels of the biosynthetic gene *pltL*. A first experiment indicated that the *pltL* transcript is strongly induced in darkness, and the difference compared to light was highest after 16 h (Supplementary Fig. 4a). An independent experiment using RNA from 100 ml of culture, which was necessary due to the low cell density in this early growth phase, confirmed that *pltL* is induced more than 100-fold after 16 h in the dark compared to light (Fig. 4a). As a complementary tool to study regulation at the transcriptional level, a *P. protegens* reporter strain with a transcriptional fusion of *gfp* to the *pltL* promoter (*P*_*pltL*_*::gfp*)^35^ was used. Growth of this reporter strain in TAP medium gave first hints for a higher transcriptional activity from the *pltL* promoter in darkness (Supplementary Fig. 4b). Owing to the low green fluorescence of the *P*_*pltL*_*::gfp* reporter strain in TAP medium, the reporter was additionally grown in LB medium where the fluorescence was higher (Supplementary Fig. 4c). On LB plates, blue light and white light decreased the fluorescence of the reporter strain by approximately 3-fold compared to darkness, demonstrating that the *pltL* transcript levels are regulated by the activity of the promoter (Fig. 4b). Raising the white light intensity from 40 to 150 µmol photons m^-2^ s^-1^ did not further diminish *pltL* promoter activity (Fig. 4b). In contrast to blue light, red light did not influence the reporter fluorescence (Fig. 4c). Taken together, these results demonstrate that transcription from the *pltL* promoter and pyoluteorin biosynthesis are repressed by light, and they confirm the stronger influence of blue light compared to red light (Figs. 3 and 4).

**Fig. 4.**
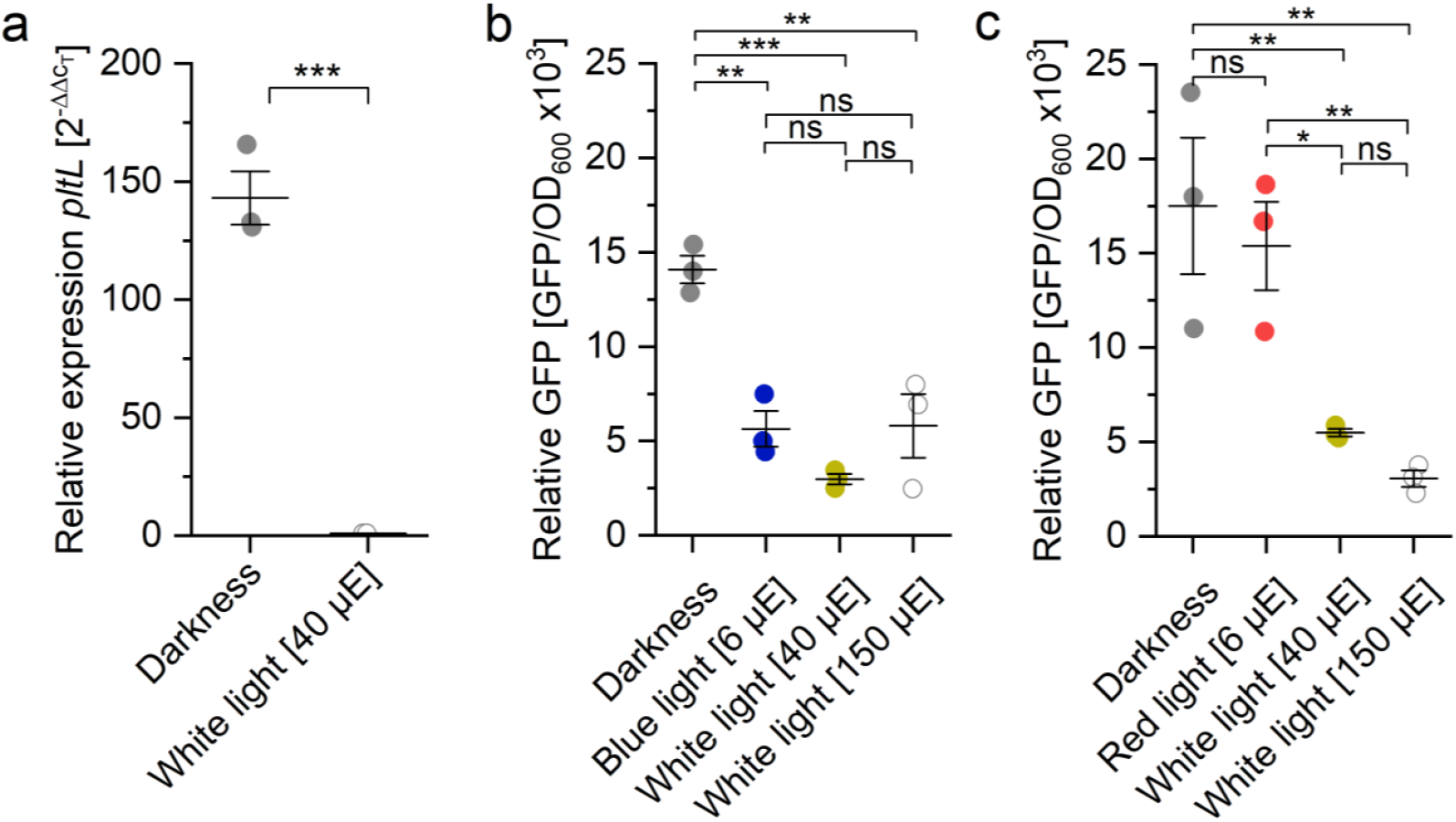
Light-dependent regulation of *pltL* transcript levels in *P. protegens*. **a** *pltL* transcript levels after 16 h in darkness or white light of 40 µmol photons m^-2^ s^-1^. *pltL* transcript levels were quantified by RT-qPCR using the *rpoD* gene for normalization. *pltL* transcript levels under white light were set to 1. A one-sample, two-tailed Student’s t-test was used for statistical analysis (***, *p* < 0.001; 3 biological replicates). **b, c** GFP expression in a *P. protegens P*_*pltL*_::*gfp* reporter strain after 48 h on LB plates under continuous darkness, or white, blue or red light (two independent datasets). The light intensities are given in µmol photons m^-2^ s^-1^. GFP fluorescence was normalized by OD_600_. A one-way ANOVA with a Tukey multiple-comparison *post-hoc* test was used for statistical analysis (*, *p* < 0.05; **, *p* < 0.01; ***, *p* < 0.001; ns, not significant; 3 biological replicates). In all three panels, the central horizontal line denotes the mean value and the whiskers denote the standard deviation.

### Evidence against a role of bacterial photoreceptors in light-regulated pyoluteorin biosynthesis

Many bacteria possess photoreceptors that regulate diverse processes^2,3^. To identify photoreceptor candidates that may repress pyoluteorin biosynthesis in the light, the proteins predicted from the *P. protegens* genome were analyzed using protein BLAST searches. Using query sequences from other *Pseudomonas* spp., we found the following predicted proteins: a LOV (light-oxygen-voltage) protein (LovP; locus tag PFL_0954), a cryptochrome/photolyase (PhrB; PFL_5147), a bacteriophytochrome (BphP; PFL_5200), and a photosensing phosphodiesterase (RmcA; PFL_5664) (Supplementary Table 3).

Knockout mutants were created by double homologous recombination to interrogate whether these possible photoreceptors regulate pyoluteorin biosynthesis. For each candidate gene, a plasmid with two selectable marker genes and homology arms for an in-frame deletion was introduced into *P. protegens* Pf-5 by biparental mating, followed by positive selection for gentamicin resistance (first homologous recombination) and negative selection against the presence of the *sacB* marker (second homologous recombination) (Supplementary Fig. 5). PCR analysis revealed that Δ*lovP*, Δ*phrB*, Δ*bphP*, and Δ*rmcA* mutants were successfully generated (Fig. 5a). The mutants were cultivated in blue light or darkness for 48 h and the pyoluteorin accumulated in the spent medium quantified by HPLC. As the wild type, the four mutants still accumulated smaller amounts of pyoluteorin in blue light compared to darkness, indicating that none of the tested putative photoreceptors regulates pyoluteorin biosynthesis (Fig. 5b). We also considered a possible redundant function of the putative photoreceptors and constructed a Δ*lovP* Δ*phrB* Δ*bphP* Δ*rmcA* quadruple mutant (Δ^*4*^), but again, this mutant still accumulated more pyoluteorin in the dark (Fig. 5b). We then made additional BLAST analyses using query sequences from other bacterial genera, resulting in the identification of a possible BLUF (blue light-sensing using flavins) protein (BluF, PFL_0280) (Supplementary Table 3). A Δ*bluF* mutant was created, which still accumulated more pyoluteorin in darkness than under blue light (Fig. 5). In summary, our results indicate that none of the five tested putative photoreceptors participates in light-repressed pyoluteorin accumulation.

**Fig. 5.**
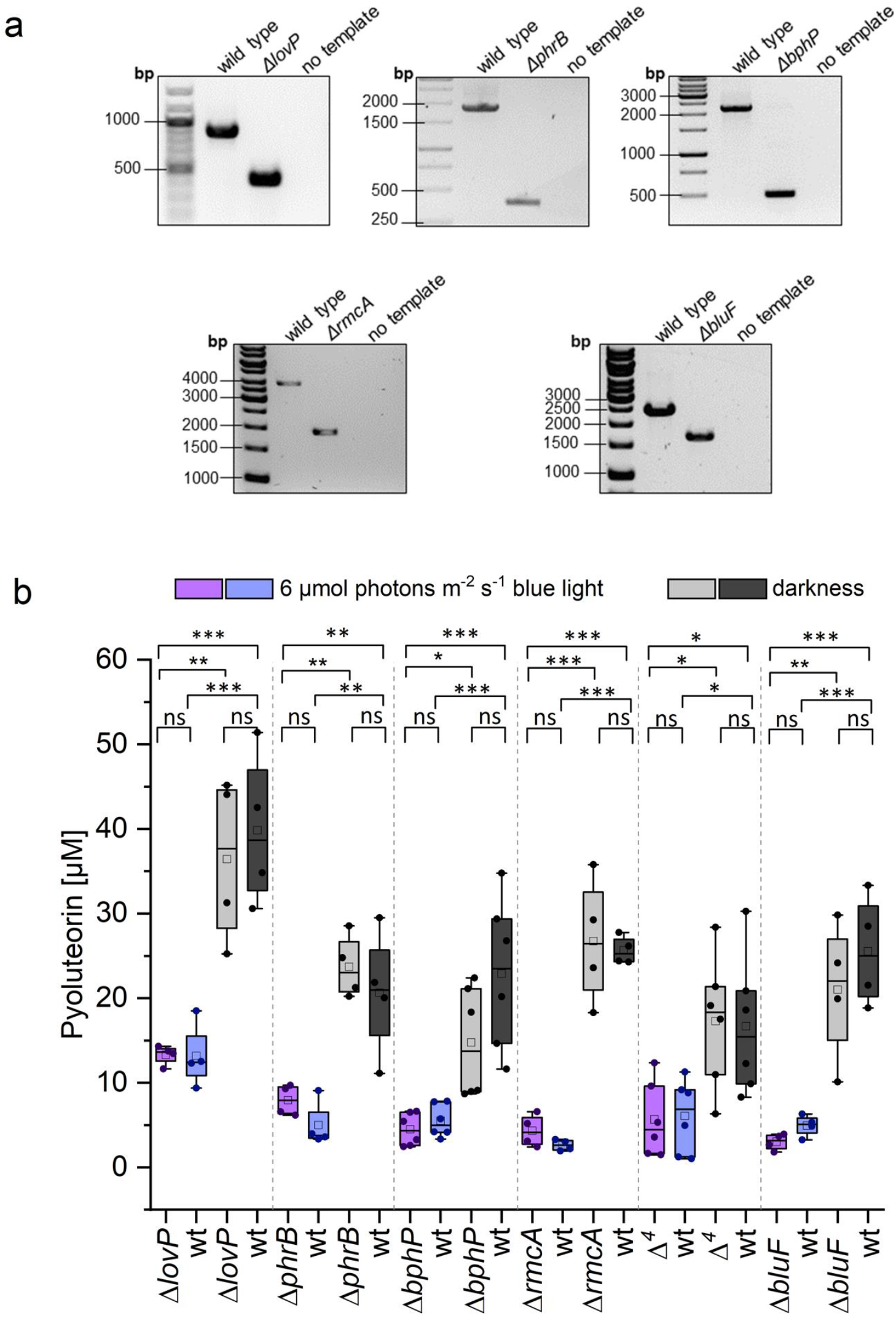
Light-repressed accumulation of pyoluteorin is unaltered in *P. protegens* mutants with disruption in five different putative photoreceptor genes. **a**, Confirmation of deletions by PCR (agarose gels). Expected product sizes: *lovP*, 872 bp (wt) vs. 425 bp (Δ*lovP*); *phrB*, 1832 bp (wt) vs. 392 bp (Δ*phrB*); *bphP*, 2222 bp (wt) vs. 496 bp (Δ*bphP*); *rmcA*, 3896 bp (wt) vs. 1903 bp (Δ*rmcA*); *bluF*, 2434 bp (wt) vs. 1657 bp (Δ*bluF*). All deletions were additionally verified by DNA sequencing. **b**, Pyoluteorin accumulation in the spent medium of mutants and wild type grown for 48 h in blue light (6 µmol photons m^-2^ s^-1^) or darkness. The spent medium of the cultures was extracted with ethyl acetate and pyoluteorin quantified via HPLC. Δ^*4*^ is a Δ*lovP* Δ*phrB* Δ*bphP* Δ*rmcA* quadruple mutant. Filled circles represent data points and open squares represent mean values. The boxes of the box plots denote the interquartile range, the central horizontal line denotes the median, and the whiskers denote the minimal and maximal values. The measurements with Δ*lovP*, Δ*phrB*, Δ*rmcA* and Δ*bluF* were performed two times independently, with two biological replicates each. The measurements with Δ*bphP* and Δ^*4*^ were performed three times independently, with two biological replicates each. A one-way ANOVA with a Tukey multiple-comparison *post-hoc* test was used for statistical analysis (*, *p* < 0.05; **, *p* < 0.01; ***, *p* < 0.001; ns, not significant).

### Light instability of PG-Cl_2_ mediates repression of pyoluteorin biosynthesis in the light

Expression of pyoluteorin biosynthetic genes and its production are activated by PG-Cl and PG-Cl_2_ that are signals converted by the enzyme PltM from PG, a compound made by PhlD of the DAPG biosynthesis pathway^16^ (Fig. 1). To test whether PG participates in the light-dependent repression of pyoluteorin biosynthesis, a *P. protegens* Δ*phlD* mutant that lacks PG was used. Strikingly, light abolished pyoluteorin biosynthesis in the Δ*phlD* mutant in the presence of exogenous PG, indicating that light interferes with PG’s role in the induction of pyoluteorin biosynthesis (Fig. 6a). To verify this result, we used *P. fluorescens* SBW25/pMRL-gfp which lacks the DAPG and pyoluteorin gene clusters but harbors the reporter plasmid pMRL-gfp containing *pltM* and *pltR* from *P. protegens* and the *pltL* promoter fused with a *gfp* gene^16^. In the presence of exogenous PG, the *pltL* promoter was more active in darkness compared to light (Fig. 6b). This finding in the heterologous host confirms the result with the Δ*phlD* mutant and demonstrates that PG, PltM and/or PltR are sufficient for the sought-after regulatory mechanism. To pinpoint the factors required for the light-dark regulation of pyoluteorin, a *P. protegens* Δ*pltM* mutant which lacks PG-Cl and PG-Cl_2_ was used. Again, light abolished pyoluteorin biosynthesis of the Δ*pltM* mutant in the presence of exogenous PG-Cl_2_ (Fig. 6a). Taken together, these results show that the light-dependent regulation of pyoluteorin accumulation only requires the signal PG-Cl_2_, which is biosynthesized via PhlD and PltM, together with its cognate transcriptional activator PltR.

**Fig. 6.**
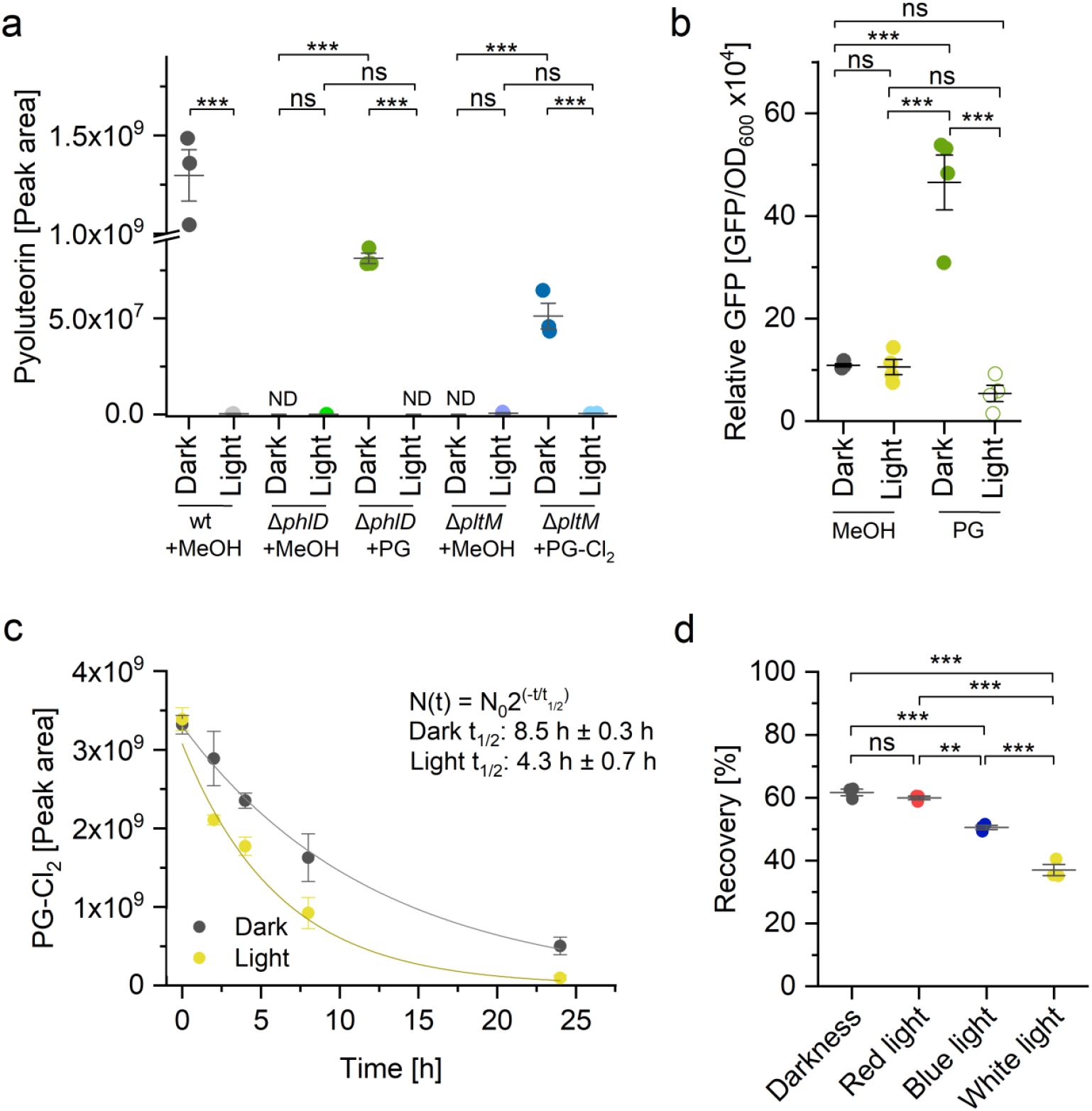
Dark-induced accumulation of pyoluteorin is regulated by the light instability of PG-Cl_2_. **a**, Pyoluteorin accumulation in the spent medium of *P. protegens* wild type and mutants grown for 24 h in darkness or white light (40 µmol photons m^-2^ s^-1^) with or without addition of 10 nM PG or PG-Cl_2_. Methanol was used as a negative control. Pyoluteorin was quantified in the spent medium via LC-MS. ND, not detectable. **b**, GFP expression in the *P. fluorescens* SBW25/pMRL-gfp reporter strain after 26 h on Nutrient Agar under continuous darkness or white light (10 µmol photons m^-2^ s^-1^) with the addition of 126 ng PG or methanol as a negative control. GFP fluorescence was normalized by OD_600_. While SBW25 lacks the *plt* gene cluster, pMRL-gfp contains *pltM, pltR* and a P_*pltL*_::*gfp* fusion^16^. **c**, *In vitro* half-lives (*t*_1/2_) of PG-Cl_2_ in darkness or white light (40 µmol photons m^-2^ s^-1^). 0.2 mM PG-Cl_2_ were incubated in 25 mM potassium phosphate, pH 7.5, at 20 °C, and PG-Cl_2_ quantified at different times by LC-MS (3 technical replicates). **d**, *In vitro* stability of of PG-Cl_2_ in darkness, red, blue or white light over a period of 6 h. The light intensities were 6 µmol photons m^-2^ s^-1^ (blue light and red light) or 40 µmol photons m^-2^ s^-1^ (white light). 0.2 mM PG-Cl_2_ were incubated in 25 mM potassium phosphate, pH 7.5, at 20 °C, and PG-Cl_2_ quantified after 0 h and 6 h (3 technical replicates). Recovery (%) is the ratio of the LC-MS signals after 6 h and 0 h. In **a, b** and **d**, filled circles represent data points, the central horizontal line denotes the mean value and the whiskers denote the standard deviation. A one-way ANOVA with a Tukey multiple-comparison *post-hoc* test was used for statistical analysis (*, *p* < 0.05; **, *p* < 0.01; ***, *p* < 0.001; ns, not significant; 3 biological replicates). In **d**, filled circles denote mean values and the whiskers denote the standard deviation.

We hypothesized that light affects the signaling function of PG-Cl_2_. To test this hypothesis, the light stability of PG-Cl_2_ was assessed in potassium phosphate buffer at pH 7.5. The results show that PG-Cl_2_ degrades faster in the light than in the dark, with half-lives of 4.3 h and 8.5 h, respectively (Fig. 6c). We did not identify any degradation products. The degradation of PG-Cl_2_ *in vitro* was faster under blue light compared to red light (Fig. 6d), which is in agreement with the differential accumulation of pyoluteorin in the spent medium of *P. protegens* under these two light qualities (Fig. 3a,b). In summary, these results demonstrate that a slower degradation of the activating signal PG-Cl_2_ in the dark leads to a higher expression of the *plt* gene cluster and a greater accumulation of pyoluteorin.

## Discussion

Among the large and diverse genus *Pseudomonas, P. protegens* Pf-5 is a prolific source of secondary metabolites and a model for a biocontrol bacterium^9^. We discovered that one of these secondary metabolites, pyoluteorin, is massively accumulated in darkness compared to light, with increases ranging from 14-fold (Fig. 3b,c) to 140-fold (Fig. 2b) and 4700-fold (wild type in Fig. 6b). While the strong increase in darkness is consistent in all experiments, the quantitative differences are likely due to different growth conditions, sampling times and analysis methods as well as the observation that pyoluteorin accumulation is a highly dynamic process (Supplementary Fig. 1f). The accumulation of pyoluteorin in the spent medium is regulated by a 140-fold induction of the *pltL* transcript (Fig. 4a). Following evidence against a role of photoreceptors in the light-dark regulation of pyoluteorin biosynthesis (Fig. 5), additional experiments with a reporter strain and mutants pointed to a role of the activating signal PG-Cl_2_ and its cognate transcription factor PltR (Fig. 6a,b). The relatively rapid degradation of PG-Cl_2_ within hours and the observation that it degrades twice as fast in the light compared to darkness (Fig. 6c) show that this activation signal is a key factor that perceives light conditions and uses this information to regulate pyoluteorin biosynthesis.

At concentrations as low as 10 nM, PG-Cl_2_ is a strong intracellular activator of pyoluteorin biosynthesis (Fig. 6b and ref. 16). In addition, it can be released by the bacteria and act as a cell-cell communication signal^16^. Our finding that a diffusible small-molecule signal perceives and transmits light information is unprecedented. In contrast to a classical chromophore, which undergoes a photochemical cycle while remaining bound to its photoreceptor^36^, PG-Cl_2_ is inactivated by degradation. PG-Cl_2_ was proposed to activate the *plt* gene cluster via binding to the transcriptional activator PltR, but the binding of PG-Cl_2_ to PltR has not been demonstrated due to unsuccessful purification of PltR (ref. 16).

In our experiments with blue light and other light qualities, the emission spectra of the blue and red-light sources seemingly did not overlap with the absorption spectrum of PG-Cl_2_ (Supplementary Figs. 6 and 7). It is thus possible that blue light and, to a lesser extent, red light promote PG-Cl_2_ degradation indirectly through an unknown mechanism. The higher energy of short-wave light may explain why blue light of 6 µmol photons m^-2^ s^-1^ inhibits pyoluteorin accumulation more strongly than the same amount of red light (Fig. 3b,c). In any case, the higher stability of PG-Cl_2_ in darkness and red light compared to blue and white light (Fig. 6d) correlates well with the differential accumulation of pyoluteorin under these conditions (Fig. 3b,c).

In addition to pyoluteorin, light and darkness influenced the accumulation of 233 other secreted metabolites in *P. protegens* (Supplementary Table 1). Differential accumulation of secreted metabolites in the spent medium may be regulated at the levels of biosynthesis, secretion, degradation, or reuptake. Given that pyoluteorin can influence the biosynthesis of several other secondary metabolites^37^, it seems likely that the differential accumulation of some metabolites in our dataset is a consequence of the different levels of pyoluteorin in light and darkness. For example, the upregulation of enantio-pyochelin (Supplementary Table 1) correlates with the higher levels of its biosynthetic enzyme PchE (ref. 37) in the presence of pyoluteorin. Therefore, the accumulation of pyoluteorin in darkness observed in this study may play an important role in the light-dependent regulation of other metabolites in *P. protegens*. Conceivably, other hits in our metabolomics dataset may involve regulation by photoreceptors and other mechanisms, and the photoreceptor mutants generated here will be valuable tools to explore this further.

In addition to *P. protegens*, pyoluteorin and *plt* gene clusters are also found in some *P. aeruginosa* strains, including a clinical isolate^38,39^. While the broad toxicity of pyoluteorin against oomycetes, fungi and bacteria^11,40^ may give these pseudomonads a fitness advantage in their habitats, its functions in nature remain largely unclear. The massive accumulation of pyoluteorin in the dark (Fig. 3b,c) suggests that pyoluteorin performs its function at night, in deeper soil layers or other light-protected environments. In light-accessible environments such as the top soil layer, the strong fluctuations of pyoluteorin may dynamically shape the composition of microbial communities during night and day. In addition to providing hints on the natural role of pyoluteorin, the elegant regulatory mechanism elucidated in this work can be leveraged for genetic engineering and optogenetics. For example, adjusting the availability of PG-Cl_2_ and light can be used to control transcription from a *pltL* promoter in genetically modified cells. Furthermore, future modification of the putative PltR/PG-Cl_2_ receptor-ligand pair may finetune the regulatory system for various applications.

## Methods

### Strains and cultivation conditions

The strains used in this study are summarized in Supplementary Table 4. Unless otherwise indicated, precultures were grown in 5 ml Tris-acetate-phosphate medium^41^ at 28 °C in darkness for 16 h with shaking at 200 rpm. The cells were washed twice with fresh TAP medium, i.e. two rounds of centrifugation (3000 × *g* for 5 min) followed by discarding the medium and resuspending the cells. The cell density was determined using a Multisizer 4e cell counter (Beckman Coulter, Krefeld, Germany) equipped with a capillary with a diameter of 30 µm. For main cultures, 100 ml of TAP medium in a 300-ml Erlenmeyer flask were inoculated with an initial density of 1×10^7^ cells ml^-1^. Cultures were incubated on a shaker at 150 rpm and 20 °C and illuminated by fluorescent lamps (Lumilux, Osram, Munich, Germany) with white light of 40 µmol photons m^-2^ s^-1^ or by LEDs (CLF floral LED modules, Plant Climatics, Wertingen, Germany) with blue or red light of 6 µmol photons m^-2^ s^-1^. The intensity of the blue light irradiation was chosen to correspond to the intensity of the white light-emitting fluorescent lamps in the range between 400 and 520 nm. Spectra of the different lamp types are shown in Supplementary Fig. 6. Erlenmeyer flasks of cultures grown in darkness were wrapped in aluminium foil.

Possible mutations in the GacS/GacA system were visualized on skim milk agar (SMA)^26^. To prepare SMA plates, 100 ml of water containing 5 g of skim milk and 1.5 of agar were autoclaved for 15 min. A culture of *P. protegens* with an initial cell density of 10^7^ cells ml^-1^ in 33 ml of TAP medium was grown at 20 °C under shaking (150 rpm) in continuous darkness or white light of 50 µmol photons m^-2^ s^-1^. After 48 h and 72 h, suitable dilutions were plated onto SMA plates, the plates incubated at 28 °C in the dark for 48 h before halo formation was assessed.

### BLASTP analyses and construction of *P. protegens* knockout mutants

Proteins predicted from the genome sequence of *P. protegens* Pf-5 (ref. 8) were searched for homologs of known bacterial photoreceptors using BLASTP on the Pseudomonas Genome Database^42^ (Supplementary Table 3). InterPro^43^ was used to predict protein domains (Supplementary Data 1). An established procedure^44^ was adapted to construct deletion mutants (Supplementary Fig. 5). Target genes with a size of up to 2000 bp were deleted completely; larger genes were partially deleted by ensuring that the region encoding the predicted chromophore-binding domain was removed (Supplementary Data 2). Left and right homology arms (LA and RA) of 750 bp each were amplified by PCR using Q5 DNA Polymerase (NEB, Ipswich, MA, USA) and the PCR products were purified with the GenElute™ PCR Clean-Up Kit (Sigma-Aldrich, St. Louis, MO, USA). Plasmid pEXG2 (ref. 45) was linearized using the restriction enzymes EcoRI and HindIII and the resulting plasmid backbone combined with the LA and RA by Gibson Assembly using the NEBuilder HIFI DNA Assembly Master Mix (NEB). After transformation of chemically competent *Escherichia coli* TOP10 (Thermo Fisher Scientific), colonies were screened for the correct plasmid by PCR using DreamTaq DNA Polymerase (Thermo Fisher Scientific); promising candidates were verified by PCR using Q5 DNA Polymerase and Sanger sequencing. All primer sequences and plasmids are listed in Supplementary Tables 5 and 6, respectively.

The verified plasmid was isolated using the GenElute Plasmid Miniprep Kit (Sigma-Aldrich) and transformed in chemically competent *E. coli* S17-1λ*pir* (ref. 46). Precultures of the donor strain S17-1λ*pir* and the acceptor strain *P. protegens* Pf-5 were grown in LB medium for 16 h at 36 °C or 28 °C, respectively. Main cultures were inoculated with an optical density at 600 nm (OD_600_) of 0.1. When the cultures reached an OD_600_ of 0.6, S17λ-1*pir* and Pf-5 were combined in ratios of 1:1, 1:2 or 1:3, centrifuged at 6000 x *g* for 2 min, the supernatant discarded and the cells resuspended in 1 ml of H_2_O. This washing procedure was repeated three more times, with the cells being resuspended in 60 µl of H_2_O the third time. 30 µl of cell suspension were applied to LB plates and the plates incubated at 28 °C for 24 h. The cells were then collected, resuspended in 200 µl of LB medium and an aliquot of the suspension spread on an LB plate containing 100 μg ml^-1^ ampicillin and 40 μg ml^-1^ gentamicin. After 24 h incubation at 28 °C, single colonies of transconjugants were restreaked on 2% agar plates containing 10% sucrose, 0.5% yeast extract and 1% tryptone and the plates incubated at 28 °C for 24 h. The deletion in the target gene was examined by colony PCR using DreamTaq DNA Polymerase; the result was verified by PCR using Q5 DNA Polymerase and Sanger sequencing of a fragment containing LA, RA and the borders on each side.

### Reverse transcription-quantitative real-time PCR

To quantify the *pltL* transcript by reverse transcription-quantitative real-time PCR (RT-qPCR), a *P. protegens* preculture was grown under continuous illumination at 40 µmol photons m^-2^ s^-1^ and 28 °C for 16 h. For main cultures, 100 ml of TAP medium were inoculated with an initial cell density of 1 × 10^7^ cell ml^-1^ and the cultures grown on a shaker at 150 rpm and 20 °C for 16 h. The complete culture was centrifuged at 3,000 × *g* and 4 °C for 5 min, the supernatant discarded and the cell pellet frozen in liquid nitrogen and stored at -80 °C until further use. The cell pellet was thawed for 2 min at room temperature and vigorously mixed for 20 s with 200 µl TE buffer (10 mM Tris-Cl, pH 8.0, 1 mM EDTA) containing 10 mg ml^-1^ lysozyme. RNA was extracted using the RNeasy Mini Kit (Qiagen, Hilden, Germany). 700 μl RTL buffer with 40 mM dithiothreitol (DTT) were added to the lysed cells and the suspension mixed vigorously for 1 min. After addition of 500 μl of ethanol, the suspension was transferred into an RNeasy Mini spin column. The column was centrifuged for 30 s at 8,000 × *g* followed by washing of the bound RNA with 350 μl RW1 buffer. 80 µl DNase I solution (10 µl DNase I + 70 µl buffer, Qiagen) were then applied to the membrane and the column incubated for 15 min at room temperature. After centrifugation, the column was washed once with 350 μl RW1 buffer and twice with 500 μl RPE buffer. The RNA was eluted with 40 μl RNase-free water and the RNA concentration quantified at 260 nm using a NanoDrop device. 4 μg of RNA were retreated with DNase I (Thermo Fisher Scientific) at 37 °C for 30 min according to the manufacturer’s instructions. After inactivation with EDTA at 65 °C for 10 min, RNA quality was assessed by agarose gel electrophoresis. 1 µg of RNA was reverse-transcribed using the SuperScript III Kit (Invitrogen, Carlsbad, CA, USA) according to the manufacturer’s manual.

For quantitative real-time PCR, a 20 µl reaction mixture contained 50 ng of cDNA, 1x DreamTaq buffer, 0.2 mM dNTPs, 0.4 µM of each primer, 2 mM MgCl_2_, 1x EvaGreen fluorescence stain (Jena BioScience, Jena, Germany), 5% DMSO and 1.25 U DreamTaq DNA Polymerase. Primer sequences are provided in Supplementary Table 5. Control samples without reverse transcriptase were included to check for possible contamination with genomic DNA. The PCR program consisted of 10 min at 95 °C, followed by 40 PCR cycles (30 s at 95 °C, 30 s at 60 °C, 30 s at 72 °C, 5 s at 82 °C for 5, quantification of fluorescence). PCR was performed in three technical replicates per biological replicate. Relative changes in *pltL* transcript levels were quantified using the 2^-ΔΔC^_T_ method^47^, with the *rpoD* gene used for normalization. The formation of specific PCR products was verified by agarose gel electrophoresis. Standard curves with different cDNA dilutions were used to ensure that the amplification efficiencies were between 90% and 110%.

### Reporter strains and quantification of GFP fluorescence

The *P. protegens*/ppltL-gfp reporter strain^35^ was grown at 28 °C under continuous white light of 40 µmol photons m^-2^s^-1^ white light in TAP or LB medium for 16 h. The culture was washed twice with fresh LB medium. For main cultures, 20 ml of medium were inoculated with an initial cell density of 10^7^ cells ml^-1^. The cultures were grown at 20 °C and shaking at 150 rpm under the light conditions described in the Results section. 1 ml of culture was harvested and washed three times with water, and GFP fluorescence was quantified using a Spark plate reader (Tecan, Männedorf, Switzerland) with excitation at 480 nm and emission detected by a 535/20 nm band pass filter. GFP fluorescence values were normalized to the corresponding OD_600_ value. The normalized fluorescence value of the control strain containing the empty vector was subtracted to correct for background noise.

For experiments with the *P. protegens*/ppltL-gfp reporter strain grown on plates, the cell density of the preculture was adjusted to an OD_600_ of 0.1. 100 µl of the culture was spread on an LB plate. After each time point, cell material was resuspended in 1 ml of water in 96-well-plates and the fluorescence determined as described above.

The *P. fluorescens* SBW25/pMRL-gfp reporter strain^16^ was grown on Nutrient Agar plates containing 50 µg ml^-1^ kanamycin and 1% glycerol. Filter paper discs containing 126 ng of PG were placed on top of the plates. Cells from an overnight culture of *P. fluorescens* SBW25/pMRL-gfp were washed and resuspended in water to an OD_600_ of 1.0. 5 µl of the cell suspension was added to the filter discs and the cells were grown in darkness or white light (10 µmol photons m^-2^ s^-1^) for 26 h at 22 °C. Filter discs carrying the reporter bacteria were then transferred to 1 ml of sterile water and vortexed to collect the cells. 200 µl of the mixture was added to a 96-well plate where the GFP fluorescence was measured using a Spark plate reader (Tecan) as described above. An excitation wavelength of 485 nm was used to measure the GFP in this experiment. Fluorescence values were normalized and corrected for background noise as described above.

### Exometabolomics (LC-MS)

500 µl aliquots of a *P. protegens* culture grown in TAP medium was taken after various time points of incubation and storage at -80 °C until analysis. After thawing, the samples were centrifuged at 16,000 x *g* for 5 min and the supernatants analyzed using a Vanquish Horizon UHPLC (Thermo Fisher Scientific, Waltham, MA, USA) coupled to an Orbitrap Q Exactive Plus mass spectrometer (Thermo Fisher Scientific) with a heated electrospray ion source (HESI). 5 µl of sample were separated using a reversed-phase column (Kinetex 1.7 µm C18 100 Å, 100 x 2.1 mm; H22-313821, Phenomenex, Torrance, CA, USA) at 25 °C. The samples were eluted with a binary gradient consisting of water + 0.05% formic acid (eluent A) and acetonitrile (ChemSolute, Th. Geyer, Renningen, Germany; eluent B). The mobile phase consisted of 5% B for 2 min, followed by a linear gradient to 100% B over a course of 13 min, 100% B for 7 min, a decrease to 5% B over 3 min, and 5% B for 5 min before injection of the next sample. The flow rate was 0.25 ml min^-1^. The HESI source was operated in positive mode at 3.5 kV. The capillary temperature was 300 °C and the auxiliary temperature was 347.8 °C with a skimmer of 15 V. Sheath gas, auxiliary gas and swap gas were set to 46.25, 10.75 and 2.31 arbitrary units, respectively. The S-lens RF level was set to 50. MS^1^ parameters were set as follows: resolution of 70,000 with a mass range of *m/z* 100-1500, 1 microscan, analyzer temperature of 30 °C, peak width of 15 s, automatic gain control (AGC) target of 1x10^6^ and ion injection time of 200 ms. MS^2^ was performed in a data-dependent mode with a pooled sample containing all samples of one timepoint. One full scan with the parameters described above was used to select 5 individual ions for isolation and fragmentation. The mass resolution for fragments was 17,000, with an adaptive mass range depending on the mass of the precursor ion. MS^2^ parameters were as follows: a peak width of 10 s; AGC target 1x10^5^; collision energy 30, 40 and 60 eV; isolation window 2 *m/z* and an ion injection time of 100 ms.

The data were evaluated using Compound Discoverer 3.3 SP3 (Thermo Fisher Scientific). All compounds with retention times of 0-22 min were included for data processing. Signals detected in a control sample with TAP medium were excluded from further analysis. For a peak to be detected, the following criteria had to be met in at least 5 samples: a mass tolerance of 5 ppm or less, a peak threshold of more than 1.5, and a peak intensity of more than 10^5^. Metabolites were identified by comparison of the MS^2^ spectra with the spectral libraries mzCloud, mzVault and GNPS2 with a precursor mass tolerance of 10 ppm and a fragment mass tolerance of 10 ppm. Identification by formula or mass alone (e.g. no MS^2^ was found or no MS^2^ was recorded) was done with ChemSpider and Mass Bank with a mass tolerance of 5 ppm. Five levels of identification confidence were adapted from ref. 48 as follows: Level 1 indicates that the correct structure was confirmed with a reference standard in MS^1^, MS^2^ and retention time matching. Level 2 indicates a probable structure by an unambiguous library spectrum match in mzVault or GNPS with a match higher than 90%. Level 3 describes a tentative structure found by a library match in mzVault lower than 90% or in mzCloud with a match higher than 90%, but the exact structure remains speculative. Level 4 indicates an unequivocal molecular formula without evidence for a possible structure. In level 5, an exact mass was measured but no formula could be assigned.

Standards of pyoluteorin (sc-391693) and DAPG (sc-206518) were obtained from Santa Cruz Biotechnology (Dallas, TX, USA), pyrrolnitrin (HY-133704) was obtained from MedChemExpress (Monmouth Junction, NJ, USA), phloroglucinol (24X9) was obtained from Carl Roth (Karlsruhe, Germany), and PG-Cl_2_ (BD01140825) was obtained from BLD Pharmatech (Shanghai, China). Rhizoxin S2 (≥ 95% HPLC) was purified from *Burkholderia rhizoxinica* as described^49^.

### Relative quantification of pyoluteorin and 2,4-dichlorophloroglucinol (LC-MS)

To quantify the stability of pyoluteorin (Fig. 3a), 75 µM pyoluteorin in 500 µl of TAP medium were incubated at 20 °C in darkness or white light of 40 µmol photons m^-2^ s^-1^. After 0 h and 48 h, 1 µl was analyzed via LC-MS as described above. For the experiment described in Fig. 6b, precultures and main cultures were grown as described under ‘Strains and cultivation conditions’, but 33 ml of TAP medium 100-ml Erlenmeyer flasks were used for the main cultures. 5 µl were analyzed via LC-MS as described above. To quantify the stability of PG-Cl_2_ (Fig. 6c,d), 0.2 mM PG-Cl_2_ in 25 mM potassium phosphate, pH 7.5, was incubated in an LC-MS vial at 20 °C. Immediately after LC-MS injection, the samples were returned to the respective conditions of light or darkness. 1 µl of sample were separated on the same C18 Kinetex column and with eluents A and B described above. The mobile phase consisted of 5% B for 2 min, followed by a linear gradient to 35% B over a course of 6 min, a further increase to 80% B over 20 s, 80% B for 1 min 40 s, a decrease to 5% B over 20 s, and 5% B over 1 min 40 s before injection of the next sample. The flow rate was 0.25 ml min^-1^ and the column temperature 25 °C. The mass spectrometry settings were the same as described above, but with ionization in the negative mode. To determine the half-life (t_1/2_) of PG-Cl_2_, the data were fitted to the equation N(t) = N_0_×2^(-t/t^_1/2)_, with N(t) denoting the amount of PG-Cl_2_ at different times and N_0_ the amount of PG-Cl_2_ at time zero. Curve fitting and all statistical evaluations were performed using the software OriginLab, v. 9.6.0.172.

For the targeted quantifications described in this subchapter, the peak areas of pyoluteorin or PG-Cl_2_ in the corresponding extracted ion chromatogram (expected *m*/*z* ± 5 ppm) were integrated using FreeStyleTM 1.8 Sp2 QF1 (Thermo Fisher Scientific).

### Absolute quantification of pyoluteorin (HPLC)

Precultures and main cultures were grown as described under ‘Strains and cultivation conditions’. After 48 h, the main culture was harvested by centrifugation at 8,000 × *g* and 4 °C for 25 min. The supernatant was extracted twice with 100 ml of HPLC grade ethyl acetate. The extracts were combined and the solvent removed by rotation evaporation at 40 °C. The dry residue was dissolved in 1 ml of HPLC grade methanol and the solution filtered through a 0.45 μm Ø Chromafil PET filter. The extract was then dried with gaseous nitrogen and redissolved in 200 µl of methanol. Pyoluteorin was quantified by HPLC coupled to a photodiode array detector (UltiMate 3000, Thermo Fisher Scientific) using a reversed-phase column (Nucleosil 120–5 C18, 250 × 4.6 mm; Wicom, product no. WIC 6F76I85). The HPLC conditions were previously described^14^. The absolute concentrations of pyoluteorin were quantified with the help of a standard curve (Supplementary Fig. 8). The assumption that pyoluteorin was extracted completely may lead to an underestimation of its concentration.

## Acknowledgments

We thank Prof. W. Gärtner and Dr. M. Gilbert for helpful discussions, Prof. C. Hertweck and Dr. K. Scherlach for the rhizoxin S2 standard, and Dr. K. Scherlach for helpful comments on the metabolomics data. We are grateful for funding by the Deutsche Forschungsgemeinschaft (DFG, German Research Foundation) via SFB 1127 (project ID 239748522) to MM, PS and SS, and via EXC 2051 #390713860, “Balance of the Microverse” to MM and PS; and by the Werner Siemens Stiftung (Paleobiotechnology) to PS. VH was supported by a fellowship from the International Leibniz Research School (ILRS), under the head of the Jena School for Microbial Communication. The LC-MS device was funded by the DFG (INST 268/451-1 FUGG) and the Saxon Ministry for Science, Culture and Tourism (SMWK).

## Author contributions

QY and SS designed the research; AD, MR and CS performed the research; RN, TJ, SZ and PS contributed methods; AD, RN, TJ and SS wrote the manuscript; all authors analyzed and discussed the data, edited the manuscript and approved its final version.

## Competing interests

The authors state that they have no competing interests.

## Supplementary Data

Supplementary Fig. 1: Metabolites secreted by *P. protegens* change depending on light/darkness.

Supplementary Fig. 2: Molecular networking revealed 32 rhizoxin derivatives.

Supplementary Fig. 3: Evidence against spontaneous mutations in the GacS/GacA regulatory system during growth of *P. protegens* in TAP medium in light or darkness.

Supplementary Fig. 4: Light-dependent regulation of *pltL* transcript levels in *P. protegens*.

Supplementary Fig. 5: Strategy used to delete putative photoreceptor genes in *P. protegens* by stepwise double homologous recombination.

Supplementary Fig. 6: Spectra of lamps used for all experiments.

Supplementary Fig. 7: Absorbance spectrum of 25 µM PG-Cl_2_ in 25 mM potassium phosphate, pH 7.5.

Supplementary Fig. 8: HPLC standard curve for the quantification of pyoluteorin.

Supplementary Table 1: Metabolites differentially accumulated in the spent medium of *P. protegens* in light and darkness (234 metabolites).

Supplementary Table 2: Cell densities of *P. protegens* wild type and photoreceptor knock-out mutants are not influenced by light.

Supplementary Table 3: Protein BLAST search for photoreceptors encoded in the genome of *P. protegens*.

Supplementary Table 4: List of strains used in this study

Supplementary Table 5: Primer used for in-frame deletion mutation of *P. protegens*.

Supplementary Table 6: List of plasmids used in this study

Supplementary data 1: Sequences of photoreceptor candidates from *P. protegens* with domains predicted by InterPro

Supplementary data 2: Sequences of *P. protegens* deletion mutants

## Notes

### Competing Interest Statement

The authors have declared no competing interest.

## References

1. van der Horst, M. A., Key, J. & Hellingwerf, K. J. Photosensing in chemotrophic, non-phototrophic bacteria: let there be light sensing too. Trends Microbiol. 15, 554–562 (2007).

2. Losi, A. & Gärtner, W. A light life together: photosensing in the plant microbiota. Photochem. Photobiol. Sci. 20, 451–473 (2021).

3. Santamaría-Hernando, S. et al. Pseudomonas syringae pv. tomato exploits light signals to optimize virulence and colonization of leaves. Environ. Microbiol. 20, 4261–4280 (2018).

4. Parte, A. C., Sardà Carbasse, J., Meier-Kolthoff, J. P., Reimer, L. C. & Göker, M. List of Prokaryotic names with Standing in Nomenclature (LPSN) moves to the DSMZ. Int. J. Syst. Evol. Microbiol. 70, 5607–5612 (2020).

5. Wang, L., Zhang, X., Lu, J. & Huang, L. Microbial diversity and interactions: Synergistic effects and potential applications of Pseudomonas and Bacillus consortia. Microbiol. Res. 293, 128054 (2025).

6. Ramette, A. et al. Pseudomonas protegens sp. nov., widespread plant-protecting bacteria producing the biocontrol compounds 2,4-diacetylphloroglucinol and pyoluteorin. Syst. Appl. Microbiol. 34, 180–188 (2011).

7. Howell, C. R. & Stipanovic, R. D. Control of Rhizoctonia solani on cotton seedlings with Pseudomonas fluorescens and with an antibiotic produced by the bacterium. Phytopathology 69, 480–482 (1979).

8. Paulsen, I. T. et al. Complete genome sequence of the plant commensal Pseudomonas fluorescens Pf-5. Nat. Biotechnol. 23, 873–878 (2005).

9. Loper, J. E., Kobayashi, D. Y. & Paulsen, I. T. The genomic sequence of Pseudomonas fluorescens Pf-5: insights into biological control. Phytopathology 97, 233–238 (2007).

10. Rodriguez, F. & Pfender, W. F. Antibiosis and antagonism of Sclerotinia homoeocarpa and Drechslera poae by Pseudomonas fluorescens Pf-5 in vitro and in planta. Phytopathology 87, 614–621 (1997).

11. Zhao, Q. et al. Pyoluteorin-deficient Pseudomonas protegens improves cooperation with Bacillus velezensis, biofilm formation, co-colonizing, and reshapes rhizosphere microbiome. npj Biofilms Microbiomes 10, 145 (2024).

12. Loper, J. E. et al. Rhizoxin analogs, orfamide A and chitinase production contribute to the toxicity of Pseudomonas protegens strain Pf-5 to Drosophila melanogaster. Environ. Microbiol. 18, 3509–3521 (2016).

13. Aiyar, P. et al. Antagonistic bacteria disrupt calcium homeostasis and immobilize algal cells. Nat. Commun. 8, 1756 (2017).

14. Rose, M. M. et al. The bacterium Pseudomonas protegens antagonizes the microalga Chlamydomonas reinhardtii using a blend of toxins. Environ. Microbiol. 23, 5525–5540 (2021).

15. Hotter, V. et al. A polyyne toxin produced by an antagonistic bacterium blinds and lyses a Chlamydomonad alga. Proc. Natl. Acad. Sci. U. S. A. 118, e2107695118 (2021).

16. Yan, Q., Philmus, B., Chang, J. H. & Loper, J. E. Novel mechanism of metabolic coregulation coordinates the biosynthesis of secondary metabolites in Pseudomonas protegens. eLife 6, e22835 (2017).

17. Thomas, M. G., Burkart, M. D. & Walsh, C. T. Conversion of L-proline to pyrrolyl-2-carboxyl-S-PCP during undecylprodigiosin and pyoluteorin biosynthesis. Chem. Biol. 9, 171–184 (2002).

18. Dorrestein, P. C., Yeh, E., Garneau-Tsodikova, S., Kelleher, N. L. & Walsh, C. T. Dichlorination of a pyrrolyl-S-carrier protein by FADH_2_-dependent halogenase PltA during pyoluteorin biosynthesis. Proc. Natl. Acad. Sci. U. S. A. 102, 13843–13848 (2005).

19. Kraus, J. & Loper, J. E. Characterization of a genomic region required for production of the antibiotic pyoluteorin by the biological control agent Pseudomonas fluorescens Pf-5. Appl. Environ. Microbiol. 61, 849–854 (1995).

20. Nowak-Thompson, B., Gould, S. J. & Loper, J. E. Identification and sequence analysis of the genes encoding a polyketide synthase required for pyoluteorin biosynthesis in Pseudomonas fluorescens Pf-5. Gene 204, 17–24 (1997).

21. Nowak-Thompson, B., Chaney, N., Wing, J. S., Gould, S. J. & Loper, J. E. Characterization of the pyoluteorin biosynthetic gene cluster of Pseudomonas fluorescens Pf-5. J. Bacteriol. 181, 2166–2174 (1999).

22. Yi, D. et al. A nonfunctional halogenase masquerades as an aromatizing dehydratase in biosynthesis of pyrrolic polyketides by type I polyketide synthases. ACS Chem. Biol. 17, 1351–1356 (2022).

23. Yan, Q., Liu, M., Kidarsa, T., Johnson, C. P. & Loper, J. E. Two pathway-specific transcriptional regulators, PltR and PltZ, coordinate autoinduction of pyoluteorin in Pseudomonas protegens Pf-5. Microorganisms 9, 1489 (2021).

24. Hassan, K. A. et al. Inactivation of the GacA response regulator in Pseudomonas fluorescens Pf-5 has far-reaching transcriptomic consequences. Environ. Microbiol. 12, 899–915 (2010).

25. Stallforth, P. et al. A bacterial symbiont is converted from an inedible producer of beneficial molecules into food by a single mutation in the gacA gene. Proc. Natl. Acad. Sci. U. S. A. 110, 14528–14533 (2013).

26. Song, C., Kidarsa, T. A., van de Mortel, J. E., Loper, J. E. & Raaijmakers, J. M. Living on the edge: emergence of spontaneous gac mutations in Pseudomonas protegens during swarming motility. Environ. Microbiol. 18, 3453–3465 (2016).

27. Zhang, Q. et al. Role of Vfr in the regulation of antifungal compound production by Pseudomonas fluorescens FD6. Microbiol. Res. 188-189, 106–112 (2016).

28. Youard, Z. A., Mislin, G. L. A., Majcherczyk, P. A., Schalk, I. J. & Reimmann, C. Pseudomonas fluorescens CHA0 produces enantio-pyochelin, the optical antipode of the Pseudomonas aeruginosa siderophore pyochelin. J. Biol. Chem. 282, 35546–35553 (2007).

29. Kirner, S. et al. Functions encoded by pyrrolnitrin biosynthetic genes from Pseudomonas fluorescens. J. Bacteriol. 180, 1939–1943 (1998).

30. Pesci, E. C. et al. Quinolone signaling in the cell-to-cell communication system of Pseudomonas aeruginosa. Proc. Natl. Acad. Sci. U. S. A. 96, 11229–11234 (1999).

31. Kusebauch, B., Scherlach, K., Kirchner, H., Dahse, H.-M. & Hertweck, C. Antiproliferative effects of ester- and amide-functionalized rhizoxin derivatives. Chemmedchem 6, 1998–2001 (2011).

32. Scherlach, K., Brendel, N., Ishida, K., Dahse, H.-M. & Hertweck, C. Photochemical oxazole–nitrile conversion downstream of rhizoxin biosynthesis and its impact on antimitotic activity. Org. Biomol.Chem. 10, 5756–5759 (2012).

33. Brendel, N., Partida-Martinez, L. P., Scherlach, K. & Hertweck, C. A cryptic PKS-NRPS gene locus in the plant commensal Pseudomonas fluorescens Pf-5 codes for the biosynthesis of an antimitotic rhizoxin complex. Org. Biomol.Chem. 5, 2211–2213 (2007).

34. Loper, J. E., Henkels, M. D., Shaffer, B. T., Valeriote, F. A. & Gross, H. Isolation and identification of rhizoxin analogs from Pseudomonas fluorescens Pf-5 by using a genomic mining strategy. Appl. Environ. Microbiol. 74, 3085–3093 (2008).

35. Yan, Q. et al. The rare codon AGA is involved in regulation of pyoluteorin biosynthesis in Pseudomonas protegens Pf-5. Front. Microbiol. 7, 497 (2016).

36. Morozov, D., Modi, V., Mironov, V. & Groenhof, G. The photocycle of bacteriophytochrome is initiated by counterclockwise chromophore isomerization. J. Phys. Chem. Lett. 13, 4538–4542 (2022).

37. Luo, L.-M. et al. Pyoluteorin regulates the biosynthesis of 2,4-DAPG through the TetR family transcription factor PhlH in Pseudomonas protegens Pf-5. Appl. Environ. Microbiol. 90, e01743–01723 (2024).

38. Li, S., Huang, X., Wang, G. & Xu, Y. Transcriptional activation of pyoluteorin operon mediated by the LysR-type regulator PltR bound at a 22 bp lys box in Pseudomonas aeruginosa M18. PLoS ONE 7, e39538 (2012).

39. Winstanley, C. et al. Newly introduced genomic prophage islands are critical determinants of in vivo competitiveness in the Liverpool Epidemic Strain of Pseudomonas aeruginosa. Genome Res. 19, 12–23 (2009).

40. Ohmori, T., Hagiwara, S.-i., Ueda, A., Minoda, Y. & Yamada, K. Production of pyoluteorin and its derivatives from n-paraffin by Pseudomonas aeruginosa S10B2. Agric. Biol. Chem. 42, 2031–2036 (1978).

41. Gorman, D. S. & Levine, R. P. Cytochrome f and plastocyanin: their sequence in the photosynthetic electron transport chain of Chlamydomonas reinhardi. Proc. Natl. Acad. Sci. U. S. A. 54, 1665–1669 (1965).

42. Winsor, G. L. et al. Enhanced annotations and features for comparing thousands of Pseudomonas genomes in the Pseudomonas Genome Database. Nucleic Acids Res. 44, D646–D653 (2015).

43. Blum, M. et al. InterPro: the protein sequence classification resource in 2025. Nucleic Acids Res. 53, D444–D456 (2025).

44. Zhang, S., Mukherji, R., Chowdhury, S., Reimer, L. & Stallforth, P. Lipopeptide-mediated bacterial interaction enables cooperative predator defense. Proc. Natl. Acad. Sci. U. S. A. 118, e2013759118 (2021).

45. Hmelo, L. R. et al. Precision-engineering the Pseudomonas aeruginosa genome with two-step allelic exchange. Nat. Protoc. 10, 1820–1841 (2015).

46. de Lorenzo, V. & Timmis, K. N. in Methods Enzymol. Vol. 235 386–405 (Academic Press, 1994).

47. Livak, K. J. & Schmittgen, T. D. Analysis of relative gene expression data using realtime quantitative PCR and the 2−ΔΔCT method. Methods 25, 402–408 (2001).

48. Schymanski, E. L. et al. Identifying small molecules via high resolution mass spectrometry: communicating confidence. Environ. Sci. Technol. 48, 2097–2098 (2014).

49. Scherlach, K., Partida-Martinez, L. P., Dahse, H.-M. & Hertweck, C. Antimitotic rhizoxin derivatives from a cultured bacterial endosymbiont of the rice pathogenic fungus Rhizopus microsporus. J. Am. Chem. Soc. 128, 11529–11536 (2006).

