## Supplementary Data for "A light-labile signal molecule acts as a photoregulator of secondary metabolite biosynthesis in a heterotrophic bacterium"

#### Table of contents

1. Supplementary figures 2

2. Supplementary tables 9

3. Supplementary data 15

4. Supplementary references 21

#### 1. Supplementary figures

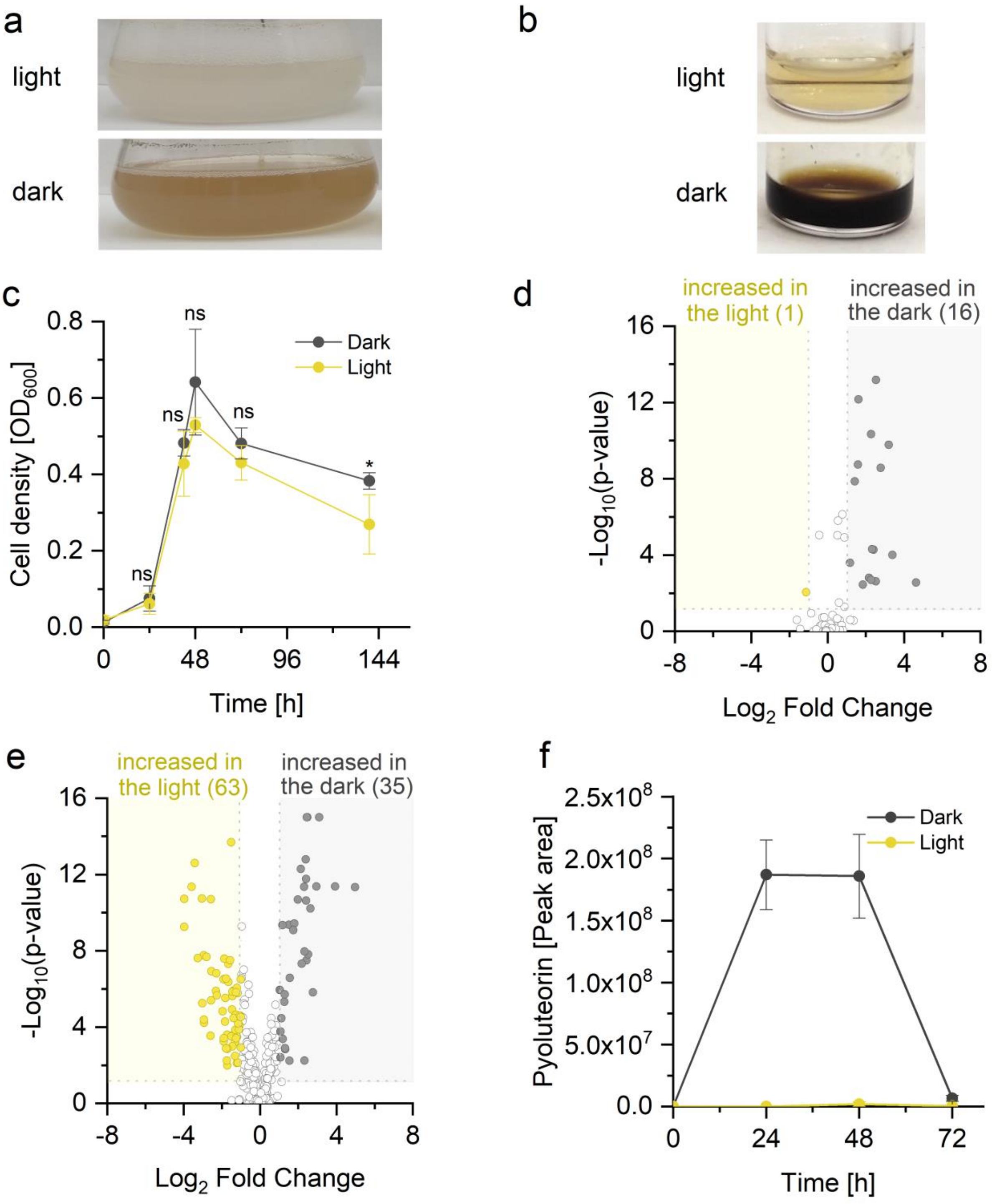


**Supplementary Fig. 1: Metabolites secreted by *P. protegens* change depending on light/darkness.** (a) Photos of *P. protegens* cultures grown under continuous light (40 µmol photons $m^{-2}$ $s^{-1}$) or darkness. (b) Extract from the spent medium of a *P. protegens* cultures grown for 48 h under 40 µmol photons $m^{-2}$ $s^{-1}$ light or in the dark. (c) Growth curves of *P. protegens* grown under continuous light (40 µmol photons $m^{-2}$ $s^{-1}$) or darkness (mean ± standard deviation). An unpaired two-tailed Student's t-test was used for statistical comparisons of the cell densities between light and darkness at each timepoint (*, $p < 0.05$; ns, not significant; 4 biological replicates). Aliquots after 0 h, 24 h, 48 h and 72 h were taken for LC-MS analyses. (d) Volcano plot depicting changes between light and darkness after 24 h (same experiment as in Fig. 2). In addition to the metabolites shown, one metabolite was only detected in the light and 7 metabolites were only detected in the dark (not shown in the volcano plot). (e) Volcano plot depicting changes between light and darkness after 72 h (same experiment as in Fig. 2). In addition to the metabolites shown in the volcano plot, 57 metabolites were only detected in the light and 57 metabolites were only detected in the dark (not shown in the volcano plot). (f) Pyoluteorin levels at different times of the metabolomics experiment in light and darkness (mean ± standard deviation).

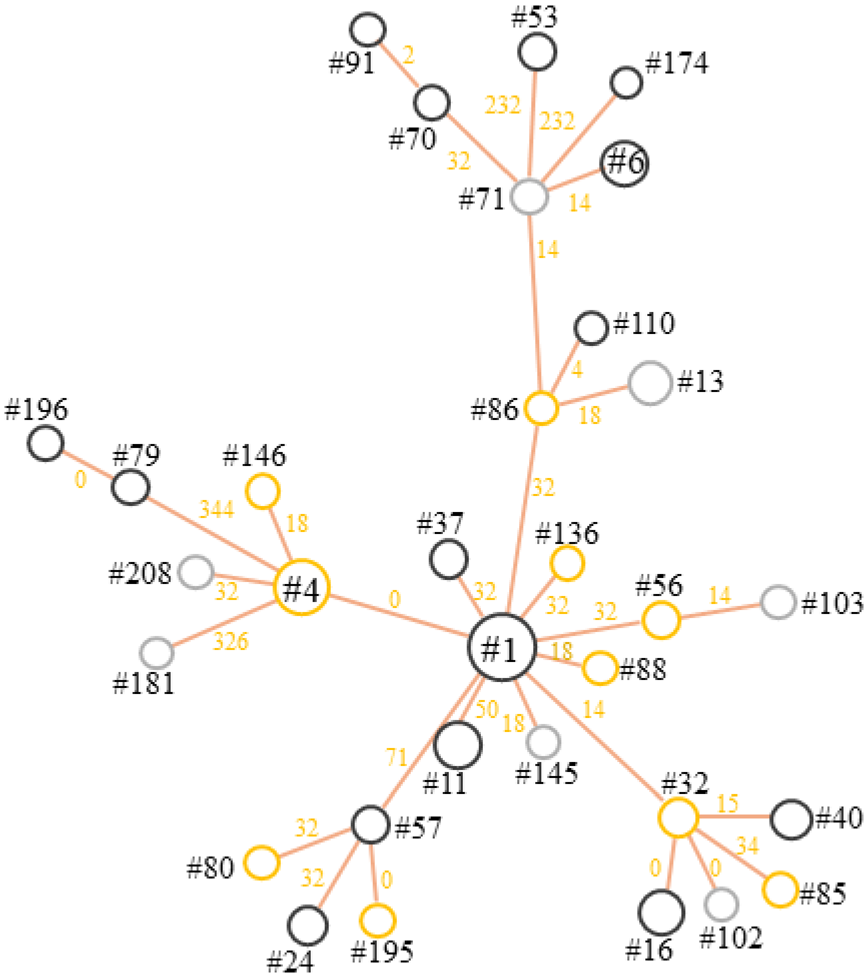


**Supplementary Fig. 2: Molecular networking revealed 32 rhizoxin derivatives.** The molecular network was generated in Compound Discoverer. The metabolites were sorted based on their similarities to rhizoxin S2 (#1). Metabolite numbers (#) are the same as in Supplementary Table 1. Metabolites were included in the network if the following criteria were met: score ≥ 50, coverage ≥ 70, and number of matched fragments ≥ 3. The size of a node provides information about the maximal amount (peak area) of a metabolite considered over the entire experiment (all timepoints and light conditions). The color of a node describes the amount of the metabolite after 48 h as follows: yellow indicates a higher abundance in the light, black indicates a higher abundance in the dark, and gray indicates no difference. The mass differences between adjacent metabolites are shown next to the branches.

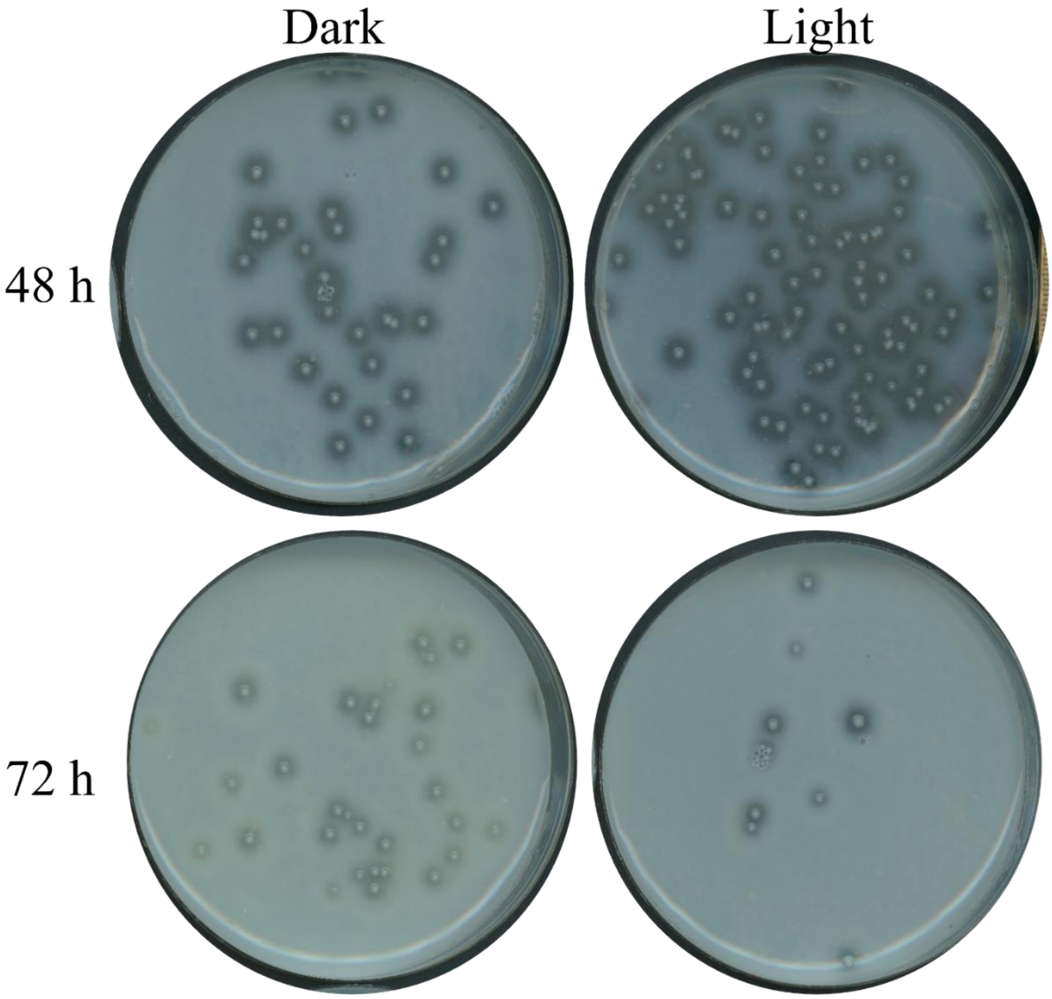


**Supplementary Fig. 3: Evidence against spontaneous mutations in the GacS/GacA regulatory system during growth of *P. protegens* in TAP medium in light or darkness.** *P. protegens* was grown in TAP medium in white light (50 µmol photons $m^{-2}$ $s^{-1}$) or darkness. After 48 h and 72 h, aliquots of the cultures were plated on skim milk agar plates. The plates were incubated at 28 °C in darkness for 48 h before they were photographed. Since the expression of extracellular proteases is regulated by GacS/GacA, halos surrounding bacterial colonies indicate a functional GacS/GacA system.

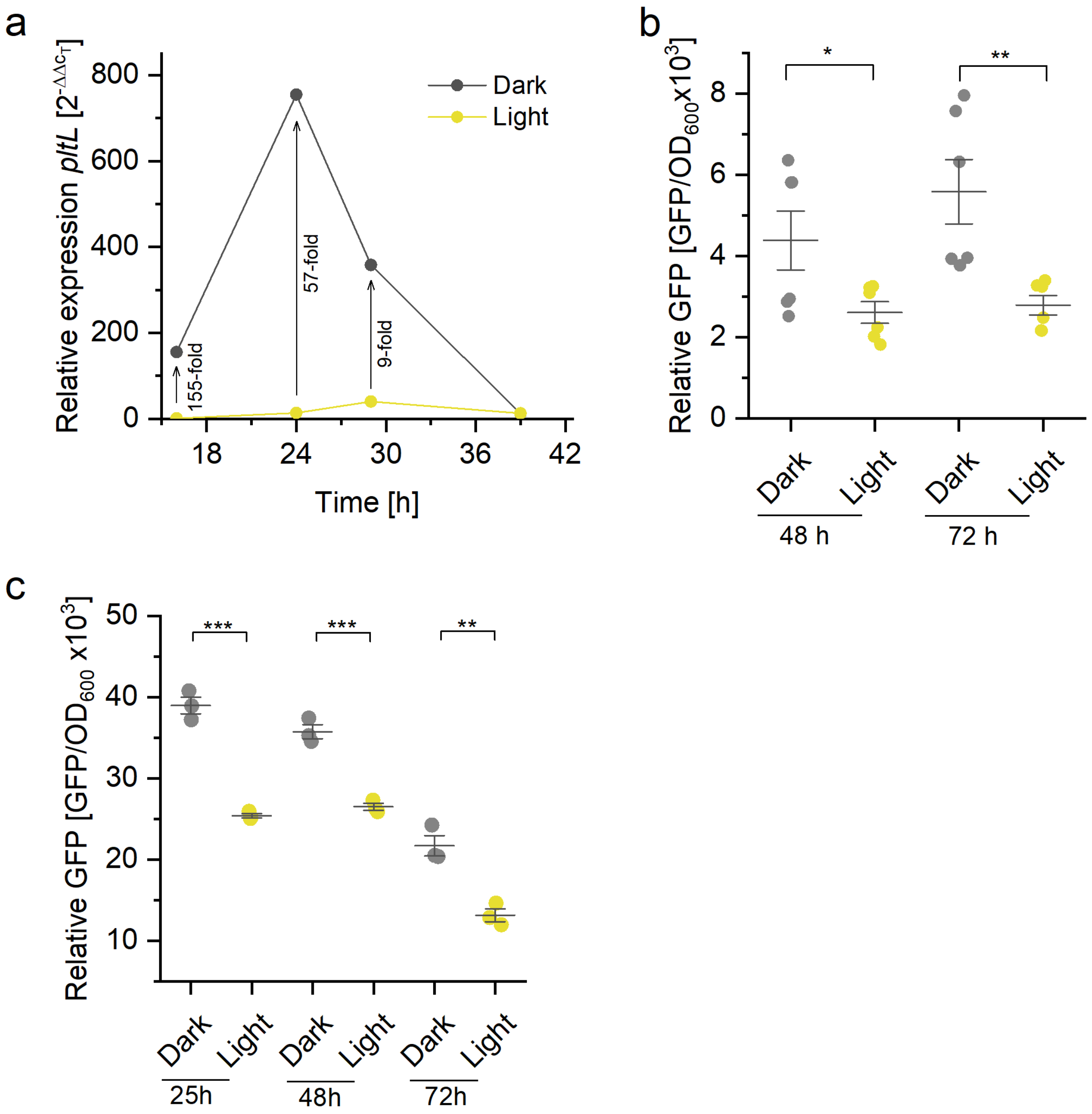


**Supplementary Fig. 4: Light-dependent regulation of *pltL* transcript levels in *P. protegens*.** (a) *pltL* transcript levels in darkness or white light of 40 µmol photons $m^{-2}$ $s^{-1}$. *pltL* transcript levels were quantified by RT-qPCR using the *rpoD* gene for normalization. The *pltL* transcript level after 16 h in the light was set to 1 (1 biological replicate with 3 technical replicates during PCR). The values next to the arrows indicate the factor by which the *pltL* transcript is upregulated in the dark compared to the light. See Supplementary Fig. 1c for growth curves recorded under the same culture conditions. (b) GFP expression in a $P_{pltL}$::*gfp* reporter strain in TAP medium in darkness and light (2 independent experiments with 3 biological replicates each). (c) GFP expression in a $P_{pltL}$::*gfp* reporter strain in LB medium in darkness and light (3 biological replicates). In (b) and (c), GFP fluorescence was normalized by $OD_{600}$. An unpaired two-tailed Student's t-test was used to compare the fluorescence values between light and darkness (*, $p < 0.05$; **, $p < 0.01$; ***, $p < 0.001$). The central horizontal line denotes the mean value and the whiskers denote the standard deviation.

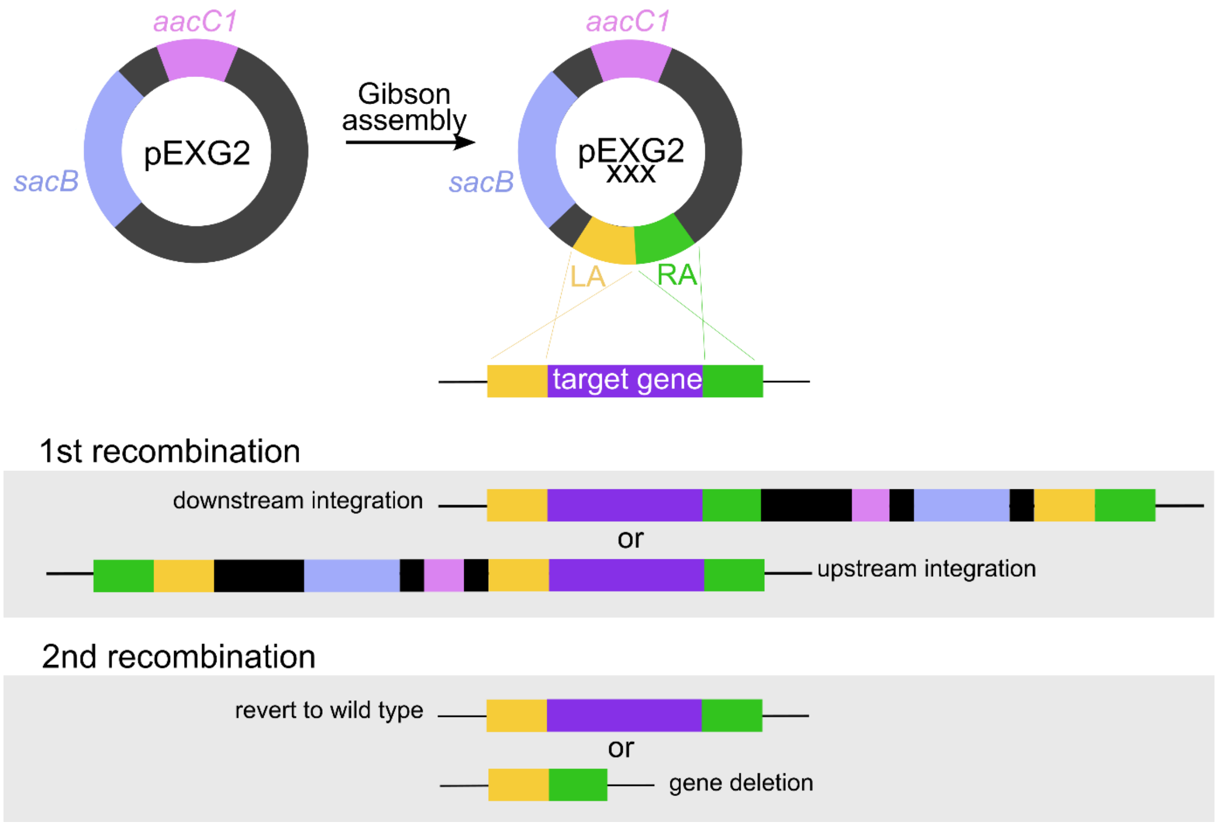


**Supplementary Fig. 5: Strategy used to delete putative photoreceptor genes in *P. protegens* by stepwise double homologous recombination.** Left and right homology arms (LA and RA) of 750 bp each were amplified by PCR and cloned into plasmid pEXG2 by Gibson assembly. After transformation of *P. protegens* by biparental mating with *E. coli* S17-1λ*pir* harboring the verified plasmid, positive selection for gentamicin resistance (*aacC1* marker) was used to select bacteria that have undergone a first homologous recombination. Ampicillin contained in the same plates additionally selected for *P. protegens* and against the *E. coli* donor strain. In the next step, negative selection on sucrose-containing plates was used to select bacteria that have undergone a second homologous recombination and eliminated the *sacB* marker. PCR was then used to identify bacteria with a successful deletion in the gene of interest.

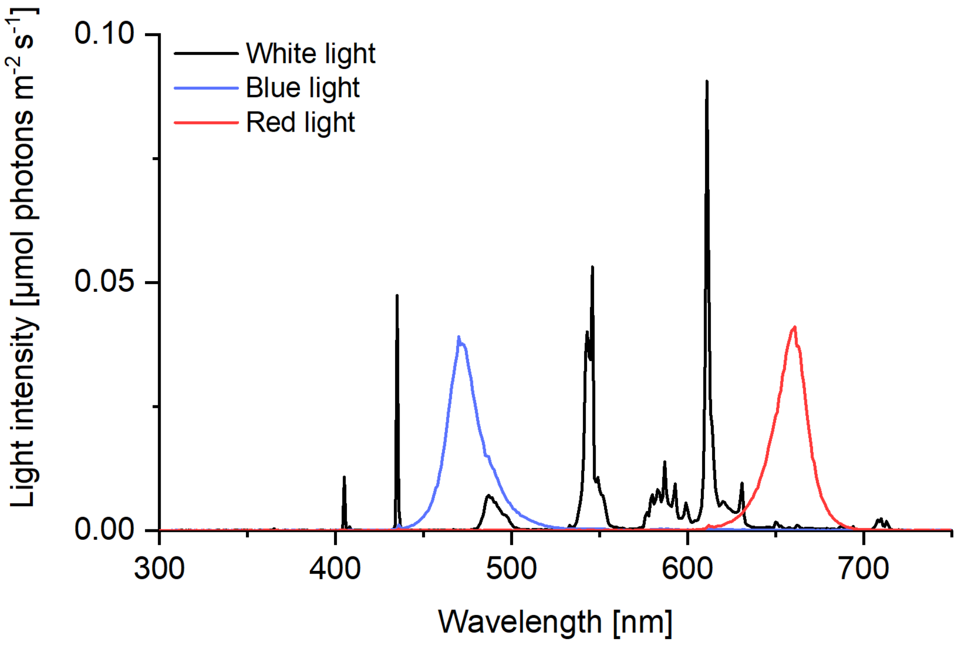


**Supplementary Fig. 6: Spectra of lamps used for all experiments.** The spectra of a white fluorescent lamp and blue and red LEDs are shown.

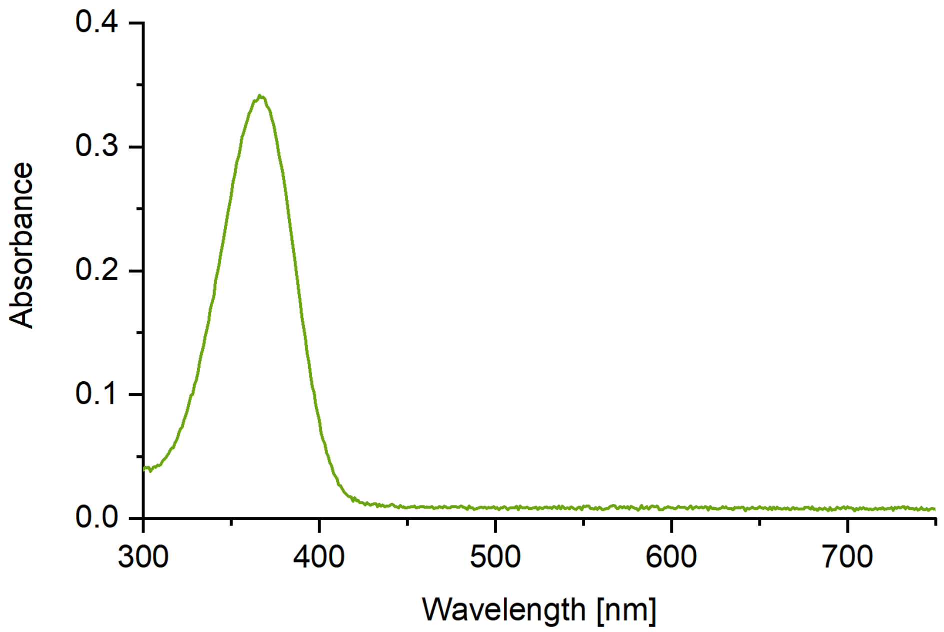


**Supplementary Fig. 7: Absorbance spectrum of 25 µM PG-$Cl_2$ in 25 mM potassium phosphate, pH 7.5.** The spectrum was measured in an M6 UV/visible spectrophotometer (VWR).

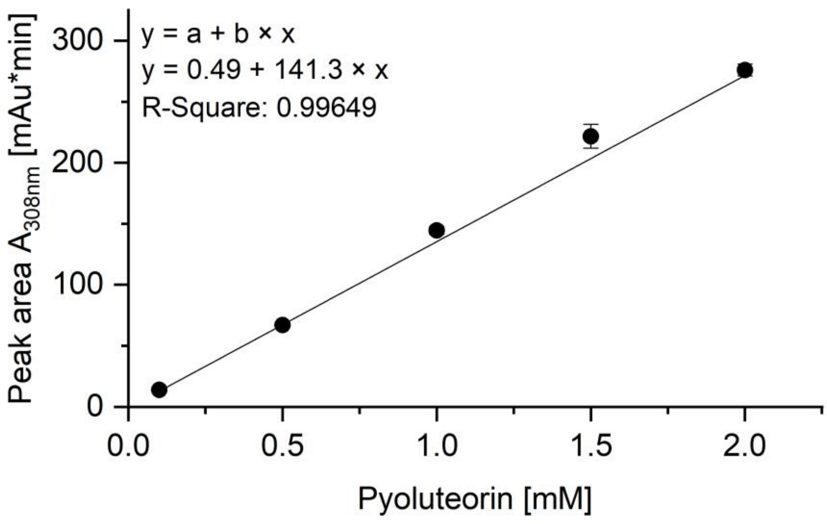


**Supplementary Fig. 8: HPLC standard curve for the quantification of pyoluteorin.** The peaks were integrated in the chromatogram recorded at 308 nm. Five different concentrations were measured in three technical replicates each. Points represent mean values, and whiskers represent standard deviations (in most cases, the standard deviation is smaller than the data point).

#### 2. Supplementary tables

**Supplementary Table 2: Cell densities of *P. protegens* wild type and photoreceptor knock-out mutants are not influenced by light.** A one-way ANOVA with a Tukey multiple-comparison *post-hoc* test was used to statistically compare the cell densities between light (blue, red or white light) and darkness, and between mutants and wild type grown under the same conditions. In all cases, there was no statistically significant difference between these cell densities ($p > 0.05$). Abbreviations: wt, wild type; SD, standard deviation

| Strain/ condition | Cell density ± SD [cells $ml^{-1}$] after 48 h | Experiment (number of replicates) |
|---|---|---|
| wt darkness | (1.92 ± 0.38) x $10^9$ | Fig. 3b (n = 6) (The cell densities were only measured in the second and third experiments.) |
| wt blue light | (2.32 ± 0.47) x $10^9$ | |
| wt white light | (1.49 ± 0.39) x $10^9$ | |
| wt darkness | (1.76 ± 0.18) x $10^9$ | Fig. 3c (n = 9) |
| wt red light | (1.88 ± 0.13) x $10^9$ | |
| wt white light | (1.90 ± 0.25) x $10^9$ | |
| Δ*lovP* blue | not measured | Fig. 5b |
| wt blue | not measured | |
| Δ*lovP* dark | not measured | |
| wt dark | not measured | |
| Δ*phrB* blue | (3.31 ± 0.33) x $10^9$ | Fig. 5b (n = 4) |
| wt blue | (3.97 ± 1.56) x $10^9$ | |
| Δ*phrB* dark | (2.77 ± 0.46) x $10^9$ | |
| wt dark | (2.55 ± 0.68) x $10^9$ | |
| Δ*bphP* blue | (2.16 ± 0.11) x $10^9$ | Fig. 5b (n = 6) |
| wt blue | (2.31 ± 0.33) x $10^9$ | |
| Δ*bphP* dark | (2.13 ± 0.16) x $10^9$ | |
| wt dark | (2.03 ± 0.19) x $10^9$ | |
| Δ*bluF* blue | (2.34 ± 0.28) x $10^9$ | Fig. 5b (n = 4) |
| wt blue | (1.98 ± 0.54) x $10^9$ | |
| Δ*bluF* dark | (2.30 ± 0.41) x $10^9$ | |
| wt dark | (2.20 ± 0.14) x $10^9$ | |
| Δ*rmcA* blue | (5.84 ± 0.31) x $10^8$ | Fig. 5b (n = 4) |
| wt blue | (8.66 ± 0.37) x $10^8$ | |
| Δ*rmcA* dark | (1.23 ± 0.74) x $10^9$ | |
| wt dark | (1.15 ± 0.54) x $10^9$ | |
| $\Delta^4$ blue | (1.58 ± 0.60) x $10^9$ | Fig. 5b (n = 6) |
| wt blue | (1.39 ± 0.63) x $10^9$ | |
| $\Delta^4$ dark | (1.42 ± 0.44) x $10^9$ | |
| wt dark | (1.32 ± 0.50) x $10^9$ | |

**Supplementary Table 3: Protein BLAST search for photoreceptors encoded in the genome of *P. protegens*.** For each major photoreceptor known, one representative was used as a query.[a)]

| Photoreceptor type | Known photoreceptor used as query (UniProtKB no.) | Best BLAST hit in *P. protegens* (locus identifier) | Sequence identity (%) | E value | Predicted protein domain (InterPro) | | | Typical chromophore [b)] |
|---|---|---|---|---|---|---|---|---|
| | | | | | Chromophore-binding domain | Effector domain | Additional domains | |
| LOV protein | LOV-HK from *Pseudomonas syringae* pv. *tomato* (Q881J7) (Wu *et al.*, 2013) | LovP (PFL_0954) | 45 | $9.4 \times 10^{-25}$ | PAS with LOV motif | - | - | FMN |
| Cryptochrome/ photolyase | Phr from *Pseudomonas aeruginosa* PAO1 (Q9HVD2) (Kim & Sundin, 2001) | PhrB (PFL_5147) | 67 | 0.0 | FAD-binding domain | DNA photolyase | - | FAD |
| Bacteriophytochrome | BphP1 from *Pseudomonas syringae* pv. *tomato* (Q882H5) (Wu *et al.*, 2013) | BphP (PFL_5200) | 52 | 0.0 | GAF | Histidin kinase | Phytochrome; PAS | Biliverdin |
| Photosensing phosphodiesterase | RmcA from *Pseudomonas aeruginosa* PA14 (Q02TI8) (Okegbe *et al.*, 2017) | RmcA (PFL_5664) | 66 | 0.0 | 4 PAS domains | GGDEF; EAL | - | FAD |
| BLUF protein | BlrP1 from *Klebsiella pneumoniae* (A6T8V8) (Winkler *et al.*, 2014) | BluF (PFL_0280) | 37 | 0.0 | Unknown | EAL | - | FAD |
| Rhodopsin | Bacteriorhodopsin from *Halobacterium salinarum* (P02945) (Dunn *et al.*, 1981) | - | - | - | - | - | - | Retinal |
| Photoactive yellow protein | PYP from *Halorhodospira halophila* (P16113) (Ihee *et al.*, 2005) | - | - | - | - | - | - | *p*-coumaric acid |

[a)] Abbreviations: EAL, diguanylate phosphodiesterase; FAD, flavin adenine dinucleotide; FMN, flavin mononucleotide; GAF, cGMP-specific phosphodiesterase; GGDEF, diguanylate cyclase; LOV, light-oxygen-voltage; PAS, Per-Arndt-Sim

[b)] based on Gomelsky & Hoff, 2011; Okegbe *et al.*, 2017; Losi & Gärtner, 2021

**Supplementary Table 4: List of strains used in this study**

| Strain | Genotype, description | Reference |
|---|---|---|
| *Pseudomonas protegens* strains: | | |
| Pf-5 | Wild type | Howell & Stipanovic, 1979 |
| LK270 | Δ*pltM* | Yan *et al.*, 2017 |
| JL4804 | Δ*phlD* | Quecine *et al.*, 2016 |
| PSB0001 | Δ*lovP* (deletion of putative photoreceptor gene) | This study |
| PSB0003 | Δ*phrB* (deletion of putative photoreceptor gene) | This study |
| PSB0005 | Δ*bphP* (deletion of putative photoreceptor gene) | This study |
| PSB0010 | Δ*bluF* (deletion of putative photoreceptor gene) | This study |
| PSB0031 | Δ*rmcA* (deletion of putative photoreceptor gene) | This study |
| PSB0037 | Δ*lovP* Δ*phrB* Δ*bphP* Δ*rmcA* ($\Delta^4$) | This study |
| *Pseudomonas fluorescens* strain: | | |
| SBW25 | Wild type, does not contain *phl* or *plt* genes | Silby *et al.*, 2009 |
| *E. coli* strains: | | |
| TOP10 | $F^-$ *mcrA* Δ(*mrr-hsdRMS-mcrBC*) ϕ80*lacZ*Δ*M15* Δ*lacX74 recA1 araD139* Δ(*ara-leu*)*7697 galU galK rpsL* ($Str^R$) *endA1 nupG* $\lambda^-$ | Thermo Fisher Scientific (product no. C404003) |
| S17-1λ*pir* | RP4-2 Tc::Mu-Km::Tn*7*, $Tp^rSm^r$, *thi pro hsdR*$^-$ *hsdM*$^+$ *recA*, λ-*pir* lysogen | de Lorenzo & Timmis, 1994 |

**Supplementary Table 5: Primer used for in-frame deletion mutation of *P. protegens*.**

| Primer name[a)] | Forward primer (5' → 3')[b)] | Reverse primer (5' → 3')[b)] | Description | Annealing temperature | |
|---|---|---|---|---|---|
| | | | | Polymerase | Temperature |
| pltL_qRT | CTGGAGTGGGGCATTCTCAA | ACTCGGCCTTTAGTTGCTCG | Amplification of ~200 bp of *pltL* for quantification of transcript level by RT-qPCR | DreamTaq | 60 °C |
| rpoD_qRT | GACGAAGAAGAAGCCGAAAG CG | AGCATTTCGGCGATGGCTTG | Amplification of ~200 bp of *rpoD*, used as reference gene in RT-qPCR | DreamTaq | 60 °C |
| lovP_LA | ggaagcataaatgtaaagcaGGTGCGGC CTGGGGTGGT | cgatttattaCATCCTGAAGTTCCGCT TTCGATCAGCAAG | Amplification of LA of *phrB* | Q5 | 68 °C |
| lovP_RA | cttcaggatgTAATAAATCGCGGAA TTCGTGGTC | ggaaattaattaaggtaccgCAGTGGCTTG CCTGGGTAAC | Amplification of RA of *phrB* | Q5 | 68 °C |
| lovP_colony | cgacttcttcagccgtgccgtaca | gcgtcagcaggtcacggatct | Amplification of ~250 bp of LA/RA to screen for deletion (colony PCR) | DreamTaq | 70 °C |
| lovP_seq | gcctgccagcaggtgggtttca | ccgagttgcccaagcgcctcaa | Amplification start: ~100 bp up-/downstream of LA/RA for sequencing of deletion | Q5 | 59 °C |
| phrB_LA | ggaagcataaatgtaaagcaAGCAGATC CTGACCATCC | gagaaagtcaCATGTCGGTTCCCGG ATC | Amplification of LA of *phrB* | Q5 | 66 °C |
| phrB_RA | aaccgacatgTGACTTTCTCGACAA CTTCGC | ggaaattaattaaggtaccgCAGGTTGGTG TTCAGCAC | Amplification of RA of *phrB* | Q5 | 72 °C |
| phrB_colony | gctgtattccggcaaggcactca | agctcacggacgttggcgtaga | Amplification of ~250 bp of LA/RA to screen for deletion (colony PCR) | DreamTaq | 65 °C |
| phrB_seq | ccggtgtcaacgcagtcacctt | cagtggcaaacaggtcaccgaact | Amplification start: ~100 bp up-/downstream of LA/RA for sequencing of deletion | Q5 | 65 °C |
| bphP_LA | ggaagcataaatgtaaagcaACGTCAGT GCAAAAAGCACAGGTAG | agcaggttgtAGCAGCACCTCGCGGC TG | Amplification of LA of *bphP* | Q5 | 70 °C |
| bphP_RA | aggtgctgctACAACCTGCTGGAA GACCTGC | ggaaattaattaaggtaccgGGTGCAGCA GGGCATGGG | Amplification of RA of *bphP* | Q5 | 68 °C |
| bphP_colony | tctatgtactggagggcgcga | tgaacttgatggcgttgccca | Amplification of ~250 bp of LA/RA to screen for deletion (colony PCR) | DreamTaq | 66 °C |

| | | | | | |
|---|---|---|---|---|---|
| bphP_seq | atctgatcctcaccgctgaacc | tggttccatcatttcgggctg | Amplification start: ~100 bp up-/downstream of LA/RA for sequencing of deletion | Q5 | 65 °C |
| blufP_LA | ggaagcataaatgtaaagcaCCCCATCATTTCGAACTG | aggagtccccTAATTGCGGATTTTCCTTATTG | Amplification of LA of *bluF* | Q5 | 70 °C |
| blufP_RA | tccgcaattaGGGGACTCCTCGCGTATC | ggaaattaattaaggtaccgTGGGGCCAGGACGACTAC | Amplification of RA of *bluF* | Q5 | 62 °C |
| blufP_colony | AGTTGGCCGTCGATCAGAGC | tcggtgattcattctgctaaccagtaa | Amplification of ~1200 bp of LA and complete RA to screen for deletion (colony PCR) | DreamTaq | 60 °C |
| blufP_seq | acagcaactgcagcgagc | atcttcagccgcagcatcg | Amplification start: ~100 bp up-/downstream of LA/RA for sequencing of deletion | Q5 | 67 °C |
| rmcA_LA | ggaagcataaatgtaaagcaCTATCAGGGCCTGGCCGC | tgggcggtcaCGCAGTTCCTGTTCCAGGG | Amplification of LA of *rmcA* | Q5 | 70 °C |
| rmcA_RA | aggaactgcgTGACCGCCCAGGCCAGCG | ggaaattaattaaggtaccgCTTCAAGGGGCAGGCTGACGATATAACC | Amplification of RA of *rmcA* | Q5 | 70 °C |
| rmcA_colony | tcagcaccatccagcaacactg | aggtcatagtcgcgcagcagtt | Amplification of ~250 bp of LA/RA to screen for deletion (colony PCR) | DreamTaq | 60 °C |
| rmcA_seq | agccggtctacactcgaaa | gcggctttcttgtttggtgg | Amplification start: ~100 bp up-/downstream of LA/RA for sequencing of deletion | Q5 | 66 °C |

[a] Primers ending with "_LA" and "_RA" were used to amplify the left and right homology arms of the gene of interest. Primers ending with "_colony" were used to amplify ~500 bp to screen mutants for the deletion. Primers ending with "_seq" were used to amplify a larger fragment starting ~100 bp outside the LA and RA, which was then used for sequencing. Primers ending with "_qRT" were used for RT-qPCR.

[b] Lowercase letters indicate overhangs used for Gibson Assembly

**Supplementary Table 6: List of plasmids used in this study**

| Plasmid | Description | Reference |
|---|---|---|
| pEXG2 | Gentamycin resistance, *sacB*, *mob*, *lacZα* | Hmelo *et al.*, 2015 |
| pEXG2Δ*lovP* | pEXG2 derivative for the construction of Δ*lovP* | This study |
| pEXG_*phr*21 | pEXG2 derivative for the construction of Δ*phrB* | This study |
| pEXG_*bphP*9 | pEXG2 derivative for the construction of Δ*bphP* | This study |
| pEXG_*PFL_5664*_PAS | pEXG2 derivative for the construction of Δ*rmcA* | This study |
| pEXG_*PFL_0280* | pEXG2 derivative for the construction of Δ*bluF* | This study |
| ppltL-gfp | pPROBE'-gfp(tagless) containing the *pltL* promoter fused to *gfp* encoding the green fluorescent protein ($P_{pltL}$::*gfp*) | Yan *et al.*, 2016 |
| pPROBE'-gfp(tagless) | pBBR1 containing promoterless *gfp*, kanamycin resistance, used as empty vector control together with ppltL-gfp | Miller *et al.*, 2000 |
| pMRL-gfp | pPROBE-NT containing *pltM*, *pltR* and the *pltL* promoter fused to *gfp* ($P_{pltL}$::*gfp*) | Yan *et al.*, 2017 |
| pPROBE-NT | pBBR1 containing promoterless *gfp*, kanamycin resistance, used as empty vector control together with pMRL-gfp | Miller *et al.*, 2000 |

### 3. Supplementary data

##### Supplementary data 1: Sequences of photoreceptor candidates from *P. protegens* with domains predicted by InterPro

<u>LovP</u>

Protein Twin LOV 1

MINAHLLQRMINASNDGIVVAEQEGEDNIVIYVNPAFERLTGYSADEVLYQDCRFLQSGDRDQPGLEVIRQALRQ
GRPCREVLRNYRKDGSHFWNELSITPVFNDSDQLTYFIGVQKDVSVQVKAQQRLLQLEQQLAEVQAELAALKATS
GH

<u>PhrB</u>

DNA photolyase Cryptochrome/photolyase FAD-binding domain

MQLIWLRSDLRLHDNTALSAAAQRGAAVAVYLLSPEQWLAHDDAPCKIDFWLRNLQSLSTALGRLNIPLLIRSAA
TWEQAPQVLLELCRQLQVQMLHFNQEYGIHESRRDAAVTRALQHAGISVQGHLDQLLFQPGSVLTKSGGYFQVFS
QFRKVCYSRLHTALPALVKAPGAQLPLSINADPLPQRVEGFPSPTQALRELWPAGDDEAQRRLEKFSDEQIHYYQ
SERDFPAKPGTSQLSPYLAAGVISPRQCLHAALRGNQGEFESGSPGAVTWINELLWREFYKHILVGYPRVSRHRA
FRPETEAVKWRHAPEELAAWQEARTGLPIIDAAMRQLLETGWMHNRLRMVVAMFLTKNLLIDWREGERFFMRHLI
DGDLAANNGGWQWSSSTGTDSAPYFRIFNPISQSQKFDSEGRFIKHWLPQLAGLNKKDIHNPAAMGGLFGVPGYP
APIVDLSRSRERALAAFKALPARMDAGVSHE

<u>BphP</u>

PAS GAF Phytochrome Two-component Histidine Kinase

MTPQDAQAFEQLLANCADEPIRSPGAIQPHGVLLTLSEPELRIQQISANVEALLGQPAAQVLGQPLEQLLGDTDG
QRIREVLQLPRLSDAPPLHLAVNGARFEGLLHRHQGVLMLELEIQLEHLQPQHLKEQTENLGRLLRRLQTAKTLN
ELYAISVSEIQAMTGYDRVLIYRFEEEGHGQVIAEATRPTMEVFNGLFFPASDIPQQARELYRSNWLRIIPNADY
QPVPLLPALRPDTQQALDLSFATLRSVSPIHCQYMKNMGVLSSMSISLLKGDQLWGLISCGNRQPLLVPHELRIA
CQTIGQVLSLQISAMEALDISRQREEKVTALASLDQAMRDTPDSVFDGLAQVPQLLLDLTQAGGVAIIEDKQLHC
FGNCPPQDEIRALHRWLQGTGQAVFASHHLANVYPPAASYQQVASGVLAMTLPKPVDNGVLWFRPEVKENINWSG
NPQKPLDLENSDAGLRLRPRTSFEIWKVEMAGISTKWSHGDRFAANDLRRSALEHDLARQVLREQQAVRARDELV
AVVSHDLRNPMTVISMLCGMMQKAFSSDGPHTSRRISSAIDTMQQAASRMNNLLEDLLDTSKIEAGRYSIAPQPL
DVSQMFEEAYSLLAPLALDKAIDISFHAEPDLRINADPERLFQVLSNLVGNAIKFTPKQGKVGVVAMSDGEQIVF
SVRDSGEGIPTDQLPFIFDRYWTMK EGNPNGTGLGLYITQGIIHAHGGKIEAHSEVGKGSEFRFSVPRASL

RmcA

Solute-binding protein family PAS PAS PAS PAS putative FAD binding GGDEF EAL

MPRLSAMLLLALLTWTATAGALTLTDEERSWLKTHPELRLGVDASWPPFEYRDEEGRYQGLAADYIHLIQERLSV
NLKPVEPASWTAVLEDAKQGKLDLLPGIMSTPERQSYMAFTRPYLDFPIVILAHEGGAQPHNLKELYGLKIAVVE
NYAPHELLRTHHPDLNLVAMPNVSSALQALATDEVDAVVGDLASSVWSLRQLKLDGLYVSGETPYRYQLAMGVPR
ENKVLVSILDKVLADLSPEEISTIQQHWVGSVLDHRTFWSDVLMYGLPGLLLLVTVLAVVIRINRRLSSEISRRV
ALEQELRSSEYHYRSLVESLSAIAWEARISDFTYSYVSPHAEALLGYPLAHWLIPGFWRNIIHPADLTRAQAFCD
HEVQAGRDHSLDYRVIAADGRCLWVRDIVSLIEHGHEPVMRGLMIDISETKHTEEALRLSEQKFASVFQQCPDIL
VIARLSDGCLLEVNKAFEEQIGLSAQSVVGQTATELNIWGIPGVGPGLLQRLQAGSIRNLEMPFRRHNGQLFTGL
ISAEPFDLDTTPALVVVVRDISQLKETQQQLQTSEEKFAKAFHASPDGLLLTRQRDGLLIEVNEGFSRITGFNSA
MSLDRSTLDLGIWVNLNERKQMLDLLQRDGFVRDFNCHIRRNDGQIRLCELSSRPLPIGGDDCMLTIARDITDRR
LMQEKLQQAATVFESTAEGVMITDTQQRISAVNRAFSEITGYSEREALGHTPRLLASGLHDSAFYAAMWHQLTLE
GHWQGEISNRRKNGELYPNWLTINAVRNRENQITHFVAVFADISSLKHAQARLDYQAHHDPLTGLPNRTLFESRL
LSALNSQQDNGSQGAVLFLDLDRFKHINDSLGHPVGDLLLKGIAVRLKEQLRDIDTVARLGGDEFIILLPGLHQA
SDADHIATKLLNCFTAPFQAGEHEFFISASIGTSLYPKDGCDVATLIKNADAAMYRSKAKGRNRVERYTRDLTAQ
ASERVALEHELRRAIERNELSLYFQPKISLSNHELVGAEALIRWHHPTFGDVPPEHFIPLAEENGMILLIGDWVL
EQACLQLNNWNRRFEDFGPLSVNLAGAQLRQPNLLGRIEQLLRDYDLQPGMLQLEITENFIMSQAEEALEVLHQL
KRLGVQLAIDDFGTGYSSLSYLKRLPLDFLKIDQSFVRGLPDDPHDVAIVRAIIALGRSMQFTVIAEGVENQEQQ
TFLTLEGCEQIQGYIVSLPLEAEEFCTTFLRIRVSDFSDSTAEKPSL

BluF

EAL

MTDFPTSLTSPGNRCEGCQQSQPLGFDFSFAYQPIVDLRDHSVFAHEALVRGVNGEGAGTVLGQVNDSNRYRFDQ
RCRTQAITLAAQLGMQSHLSINFMPNAVYRPELCIRSTLEAARAQRFPLDRLIFETLESQHVDNYRHLTNILREY
REFGFKTAIDDFGSGYSGLNLLADFQPDLIKLDMALVRDVDQDRVRQAIIRAIVTMCAELGVTVIAEGIESAGER
DFLSDCGIYLMQGYWFAKPAFKALAEVPAAAWNS

**Supplementary data 2: Sequences of *P. protegens* deletion mutants**

Color code:

```
Green:   Left homology arm (750 bp upstream of the deletion)
Yellow:  Right homology arm (750 bp downstream of the deletion)
Blue:    Start codon
Red:     Stop codon
```

*ΔlovP*

CGCTGGATGCCTTGCTGCCCGCCGATCACCCCTTGGCGATGAACGCCCAGGTACGCCTGGAAGAACTGGCGGACA
CGCCGTTCCTCCTGTACCAGCGCAGTTTCGTGCTCAACGACCGGCTGTTGCAGGCCTGCCAGCAGGTGGGTTTCA
CCCCCAGGGAAGGCGGGCGCAGCGGCCAGGCGGATTTCCTGGCGGCGCTGGTGGCCGCCGGCCAGGGCGTGGTGC
TGTTGCCCAGTGTGGTGGCCCGTGGCCTGGTGCGGCCTGGGGTGGTGCGCCTGACCTTGAAGGCGCCGGACTACC
TGCGATGGGATATCGCCTTCATCTGGCGCCAGGGCGCTTATCTGTCCAAGGCAGCGCAAGCCTGGCTGGCGCTGC
TGCGGGAACAGCCGGTCAGCCCCGCAGGGCGCTGATCAGCTCGGCCAGCCAGGGCTCGGCGTCGGTTTCCGGGGT
CACGCTCTCGCTGGCGTCCAGGCGCAGCATGGGCAGCACTTCCTGCACGCCCAGCTCGGCGAACAGCTCGCGCAT
CTGCTCGCCACCGCCACAGAAGGTATCGCCGTAGCTGGAATCCCCCAGGCCGATGACCGCTCCGGGCAACCCGCG
CCAGGCGGCGGGCAACTGATCGCGAATACTCGAATACAGCGGCAGCAGGTTGTCCGGCAACTCGCCCATGCCGGT
GGTGGAGGTCACGGCCAGAAACGCCTGCGGGGCGAAGGCCTGAAGCTCGGCCAGAGTGGCGCGCGAGTTGTGCCA
GGCCTCGAAACCTGCGGCGTTGAGGAGTTTGGCGGCATGCCGGGCGACTTCTTCAGCCGTGCCGTACACCGAGCC
GGAAAGGATGGCGACTTTCATCAATCTGATCCTGAAGCTGACTAAAAGGCTGGGATATTAACAGCTACGGAGTAA
AATCCCGCCTGTGCCGCCACCTCTGCTGCAAAGCGCTGGTGGACAGCGGGCCAGCCTCTCTTACACTCCAGCACT
TGCTGATCGAAAGCGGAACTTCAGGATGTAATAAATCGCGGAATTCGTGGTCGTAGGGATATTCCCCGCCGATCT
GTCCTCCACCCTCCGAGCAACATCATGCAGCGTGACGCCCTCCTCACCCAGGATGAACTGGACTTCATCCAGACC
ATGCAGCACAACCCGCAGCTCAATGTGCGTGATACCACCTCGAGCCTGATGGTCAATGGCGGCGCGCAGATCCGT
GACCTGCTGACGCGCCTGGCGGCCAATGAAAAGGTCACCATCCAGGCCCGTTTCGACAACCAGCAGATGACCTTC
CCGCTGCAGCTGGTGGAAGACGAATTTCATGCCCTGCACCTGCGGCTGGGGGTGCCGAGCATCTTCGAGGATGGG
CCCATGGTGCGCCCCTGGCGCCTGGCCCTGGAGCAGCCGGTGGCCCTGGAAAATGCCAGGGGCCAGCCTGGCCGC
CTGTGGGTTCGGGAGGTCTCGTTCAAGGGGATATTGGTGGAGATCCGCAACGGTACCCGGCCACCGCGGCAGTTC
GCCCAGTGGTTCAGCCCCTCGGGCTATGAACGGATTGCCCTGCACGGGCGCTTCGAGCGCCAGACCGAAGCCGGT
TTCTACGCCTACCGCCTGGATCAGAGCGACCTGGAGGAAACCGAGCGCCTGCGCCAGTTCATCCTCCAGCAGCAC
CGCCACAGCCATCCGGCACTGCACGCCTGAAGGCTAGCGCCTCAAAGAAACCCGCGCAGGCGCTGCTGCATCAAG
CGCCCCTCGTTACCCAGGCAAGCCACTGGCGAACCGACCAGGCTGTCCTGAGCCAGGTCCGCCGCGTCCCCGGCC
AGCAGCAATGGGCAATCCAGGGTCAGTGCCAGGCGCTTGAGGCGCTTGGGCAACTCGGCCGAGGGTGCGTGATTG
GAAAACACCACCAGGGCTACCGGCGCGACCTTTTCGCAGACCAGGGTCAGTTCCTCCAGCGGCACACCTGGGCCC
AGCAGGCGCACACCGATCGGGTCTTGCCCGAGCATCAGCGCCGCCACCAACAGCTC

*ΔphrB*

TACCTGCAGCCGCACCCCGAAGACCTCAGCCTGCGCAACGCGATCTGAGCCATGAAACCCTTCCCCCCAGCGGTC
GACGAAGACACCGCCCCGGATACCGACTGCCAGGAACAGGGCTGGCTGCCGATCCGCGAGGTGGCGCGCCAGACC
GGTGTCAACGCAGTCACCTTGCGGGCCTGGGAGCGCCGCTACGGGCTGATCGTGCCCCGGCGCACGCCCAAGGGC
CACCGGCTGTATTCCAGCGAGCAGGTGCAGCAGATCCTGACCATCCTCACCTGGCTCAATCGAGGGGTTGCGGTG
AGCCAGGTCAAGCCCTTGCTGGACAGCGACACCAGCTCCAGCCCCCAGGTGGAGAATGACTGGCTGCACCTGCGC
GCCGCCCTCGCCCGGGCCATCAGCCAGCTGGCCGAGCGCAAGGTCGACGATACCTTCAATCAGGCCATGGCGCTG
TACCCGCCACGGACCCTGTGCGAGCACCTGTTGATGCCGCTGCTGGCGGAACTGCAACAACGCTGGCAAGGCCAG
TTCGGCAGCCAGATGGAACAGGTGTTCTTCTACTCCTGGCTGCGCAGCAAGCTCGGCGCCCGGATCTATCACAAC
AACCGCCAAGTGCAGGGCGCCCCGCTGTTGCTGGTCAATCAGTCGGACCTGCCACTGGAGCCGCACCTGTGGCTC
ACCGCCTGGCTGGCCAGTAACGCCGACTGCCCGGTGGAGGTCTTCGACTGGCCGCTGCCCGCCGGCGAACTGGCC
CTGGCCACCGAACGCCTGCAGGCCCGGGGCCTGCTGCTGTATTCCGGCAAGGCACTCAACCCCGCCCAGTTGCCC
AGGCTGCTCAGCGGCGTGGCCTGCAAAAAATTCATCGTCGGACCCGCGGTGTGCATCCACCCCGCCCAGTGGTCC

GTATGTACCACTGAAATCGCCGATCTGTACCTGGCCCAGGACCCGATAGCGGCCTGCCAGCTACTGCTCGAGCGA
GGACTTTTCTGATCCGGGAACCGACATGTGACTTTCTCGACAACTTCGCCCGGGAGTTCGCCGCCCTCGACCGGC
ACAATCTCGACCGACTGGAGCAGCTCTACAGCGATGACATCCAGTTCACCGATCCTCTGCACGAGGTCCAGGGGC
TGGCCCAGTTGCGCGGCTACTTCGCCGAGCTCTACGCCAACGTCCGTGAGCTGCGCTTTGAATTCCACGGCAGCG
ACCTGTGCGCCGAGGGCCAGGGCTACCTGCGCTGGACCATGAGCTACCGCCACCCGCGCCTCAACGGCGGCGGCC
TGATCCAGGTGGCGGGCTGCTCGTACCTGCAATGGCGCGACGCCAGGGTCTACCGCCACCGCGACTATTTCGACG
CCGGTGCCCTGCTCTATGAACACTTGCCGCTACTCGGGGCGATGATCGCCTGGCTGAAACGGAGGCTCGGATGAA
GCCTGCCATGCCACGAAGGGTCTGGCTCACCGGTGCCAGCAGCGGCATCGGCGCAGCCCTGGCCGAAGAACTGCT
CAAGGCCGGGGCCCAGCTGGCGCTCAGCGCCCGTTCCCGCGAGCCCCTGGAGGCCCTGGCCAAGCGCTATCCGGG
CCGAGTGCTGGTGGTGCCCGGCGACCTCACCGATGCCTTGCAGGTACGCCAGATCGGCAAGCGCATCGCCCAGGC
CTGGGGCGCCCTGGACACGGTGATCCTGAATGCCGGGACCTGCGAATACATCGAGGTACAGCAGTTCGACGCAGC
GCTGATCGAACGGGTGCTGAACACCAACCTGCTGGCCGCCAGCTACTGCATCGAGGCCGCACTGCCGCTGTTGCG
CGCAGGCAATCGCCCGTACCTGGTGGGCGTGGCCAGTTCGGTGACCTGTTTGCCACTGCCCCGGGCCGGTGCCTA
TGGCGCCTCCAAGGCGGCGCTGCGCTATCTGCTGCAGTCCCTGCGTATCGATCTTGCTGCCGAAGGCATCGACGT
GACCCTGGTCAGCCCGGGTTTTGTCGACACGCCACTGACCGCACGCAACGATTTCC

*ΔbphP*

TGCCCAAGGGCGTCGAGCACAAACCCTGGGCCGAGCAGGAGGTCAAACTGCTGCTGATCGAACCCAAGGGCGTGC
TCAACACCGGCGATCAAGGTGGCGAACGCACCGCCCAGAACGACGTCTGGATCTGATCCTCACCGCTGAACCTGC
GAGACGGAAAAAGGAATTTCTGCCTCTTGAACACAGTCAGACACAGCAACACGTCATCGATCGACTACCCTTGGC
CGACCAATGGTGCGTCAAAAACTGTACGTCAGTGCAAAAAGCACAGGTAGAGTGACGCTCTTATTTTTCTGTGCG
ATGGCTCTAGTGCAGGGACTACGGCCTGGTCCCAGCTTTCGATGACGCCCATGGCGCGTCCATTACCTGTTGCGA
CCCCTGACCCCTGAATGCCCAAACCGCACCAAGACACCACCGCCCCCACCCTGCTTCAAGACCTGCGCGCCGGCA
CCGCGACCTTGCACGTGGCCCTGGAAAAGCGCCTGCCGTTCTTCTCCGAGCACCTCGATGCGGCCTGGTACCGGC
GCCTGATCCAGGCCTATTTCGGCTTCTATCAGCCCCTGGAGGCGGCCCTGCATGACAGCGGCCTGATCCCTGCCG
GCTACGACCCGGAGCAGCGCCGCAAGACCCCGGCCCTGGCCGGCGACCTGCAGGCCCTGGGCCTGGCCCCGGAGC
GCATCGACTCCCTGCCCCGCTGCGACCATCTGCCCGCACTGGCCAGCCCCGGCGCGTGCCTGGGCGCCCTCTATG
TACTGGAGGGCGCGACCCTGGGCGGGCAGATCCTGCGCCGGGAAATGGCCCAGCGCCTGGACCTGCACGGGGATA
ACGGCGGGGCTTTCCTCGACGTGTATGGCGCCGCCACCGGGCGGCGCTGGAAAGACTTTCTCGACTACCTGGGGC
GCATGCCCGAGGACGCGAGCGCACGACAACAAGTGGTCAGCGCCGCGCAATCCACATTTGCCTGTTTCGAGCACT
GGCTCGACAGCCGCGAGGTGCTGCTACAACCTGCTGGAAGACCTGCTGGACACCTCCAAAATCGAGGCCGGGCGC
TACAGCATCGCCCCGCAGCCGCTGGACGTCAGCCAGATGTTCGAAGAGGCCTATTCATTGCTGGCGCCACTGGCC
CTGGACAAGGCCATCGACATCAGCTTCCACGCCGAACCGGACCTCAGGATCAACGCCGACCCCGAGCGCCTGTTC
CAGGTGCTGTCGAACCTGGTGGGCAACGCCATCAAGTTCACCCCCAAACAGGGCAAGGTCGGGGTGGTGGCGATG
TCCGACGGCGAGCAGATCGTGTTCAGCGTGCGCGACTCCGGGGAGGGCATTCCCACCGACCAGTTGCCATTCATT
TTCGACCGCTACTGGACCATGAAGGAAGGCAACCCCAACGGCACCGGCCTGGGGCTGTACATCACCCAGGGCATC
ATCCACGCCCACGGCGGCAAGATCGAAGCCCACAGCGAAGTGGGCAAGGGCAGCGAGTTCCGCTTCAGCGTGCCG
CGCGCCAGCCTCTGACCCATAGCCCGGGGCTGGCCATTGAGCCCCGGGCGCTCATTCAGCCGCCTGGGCCACCAG
CCAGTCGCGAAACAACTGGCCGCTGGCGGACAGGCGCTTGTGCCGCGGATGCACCAGAAAATCCGCCTGCCCGGT
GCGGAACGCCCACTCGCCAACCCGCAGCAACTCGCCCTGGGCCACCAACTGGCTGGCCATGAAATCCCAGCCCAG
CCCCACGCCCATGCCCTGCTGCACCGCGCTGAACATCAGGGTCAGCTGGTTGAACCCCAGGGCCCCCGGTGGCGC
GCTGTAGTCCAGCCCGAAATGATGGAACCAGTCCCGCCAGTCGATGCAGTTCCACTCCGAGGCGTCCAGCTGGAT
CAACGGCAGTTCGGCCAGTTGCGCCAGCTCCTTGACCACAGGCAGCTCCAGGCGCCGGCTGACGATGGGGTAGAC
CATCTCGGCGAACAGCGATACCGCCGCCAGGGACGACCAGTGCCCCGCACCATAGAGAATGCCGAAGTCATGCTC
GGCCACGCTGGACTCGTCCATGTCGTTGTTGGCATGGATGCTCACGGTGATGTCCGGGTGCTGCTTGTTGAATGC
CAGCAGGCGCGGGAACAGCCAGAACTGCGCCACCGCGTGGGTGCATTGCACGGTGATGGTATTGGCGTCGCGCCC
GGCCCGCAGGCGGCTGACGCTGCGCAGCAGGGCTTCGAGCTGGGTGCTGACCTCGACGAACAGCGAGGCGCCGCT
GGCGGTGAGCTTCACCCCCCGGGCCTGGCGCTCGAACAGTTCGGTCTTGAGCGACAGCTCCAGGGCACGGATCTG
CTTGCTGACCGCGCTCTGGGTCAGGCACAGCTCGGTGGCCGCGCGGGTGAAGCTGGCGTTGCGGCCCACCGCCTC
GAAGGTGACCAGGGTTTCCAGCGGCGGCAGATGACGGGCGAAGCGGCTCACAAGGCATTCCCCAGGAGAATAGGT
AGGTGCCTATATATCCTTTGTCCATCCGGTCAGCAATGAC

*ΔrmcA*

CTCCTTCCTCGACGAGTAAGCCCAAGACCCCCAGCCCAGCTGGGGGTTTTGCTTTTTGCAGCCCATCCTTCTTTC
CTTCCCCGCCCCATAGCCGGTCTACACTCGAAACATTCCCCGTGCCATAACGAGACGGTTATGCCCAGACTGTCG
GCCATGCTCCTGCTGGCGCTGCTCACCTGGACCGCAACGGCTGGCGCGCTGACTCTCACCGACGAAGAACGTAGC
TGGCTCAAGACTCACCCGGAACTGCGCCTGGGTGTGGACGCTTCATGGCCGCCTTTCGAGTATCGGGATGAAGAA
GGCCGCTATCAGGGCCTGGCCGCGGACTACATCCACCTGATCCAGGAACGCCTGTCCGTAAACCTCAAGCCCGTC
GAGCCCGCCAGCTGGACCGCGGTGCTAGAGGACGCCAAGCAGGGCAAGCTGGACCTGCTGCCCGGGATCATGTCG
ACCCCCGAACGCCAGAGCTACATGGCCTTCACCCGGCCTTACCTGGACTTCCCCATCGTCATCCTCGCCCATGAA
GGCGGCGCGCAACCCCACAACCTCAAGGAACTGTACGGACTCAAGATAGCGGTGGTGGAAAACTACGCCCCTCAC
GAACTGTTGCGCACCCACCACCCGGACCTGAACCTGGTGGCCATGCCCAATGTCAGCTCGGCGCTGCAGGCCCTG
GCCACCGACGAAGTGGACGCCGTGGTCGGCGACCTCGCCTCCAGTGTCTGGAGCCTGCGTCAGCTCAAGCTCGAC
GGTCTCTACGTCAGCGGCGAAACCCCCTATCGCTATCAGTTGGCGATGGGCGTCCCGCGGGAAAACAAGGTTCTG
GTAAGCATTCTCGACAAGGTGCTCGCCGACCTCAGCCCCGAGGAAATCAGCACCATCCAGCAACACTGGGTGGGC
AGCGTGCTCGACCACAGGACCTTCTGGTCGGATGTCCTGATGTATGGCCTGCCCGGTCTTCTGCTGCTGGTGACC
GTGCTGGCGGTGGTGATCCGCATCAACCGCCGCTTGAGCTCGGAAATTTCCCGGCGGGTCGCCCTGGAACAGGAA
CTGCGTGACCCGCCCAGGCCAGCGAACGGGTAGCCCTGGAGCACGAGCTGCGGCGAGCCATAGAGCGCAACGAACT
GTCACTGTATTTCCAACCCAAGATCAGCCTCAGCAACCACGAGCTGGTGGGCGCCGAAGCCCTGATCCGCTGGCA
TCACCCCACTTTCGGCGATGTGCCCCCAGAGCACTTCATTCCCCTGGCCGAAGAAAATGGCATGATCCTGCTGAT
CGGCGACTGGGTACTGGAACAGGCCTGCCTGCAACTCAACAACTGGAACCGGCGCTTTGAGGACTTCGGCCCGCT
GTCGGTGAACCTCGCCGGCGCCCAGTTGCGCCAGCCCAACCTGCTGGGGCGCATCGAGCAACTGCTGCGCGACTA
TGACCTGCAGCCCGGAATGCTACAGCTGGAGATCACCGAGAATTTCATCATGAGCCAGGCCGAAGAAGCCCTGGA
AGTGCTGCATCAGCTCAAGCGCCTGGGCGTGCAACTGGCCATCGACGACTTCGGCACCGGCTACTCGTCCCTCAG
CTACCTCAAGCGCCTGCCCCTGGACTTTCTCAAGATCGACCAGTCCTTTGTCCGCGGGCTGCCGGACGACCCCCA
CGACGTGGCGATTGTCCGCGCCATCATCGCCCTGGGGCGCAGCATGCAGTTCACGGTGATTGCCGAAGGCGTGGA
AAACCAGGAGCAGCAGACCTTCCTCACCCTGGAAGGCTGCGAACAGATCCAGGGTTATATCGTCAGCCTGCCCCT
TGAAGCCGAGGAGTTCTGTACGACCTTCCTGCGGATCCGAGTTTCCGATTTTTCGGATAGCACAGCCGAGAAACC
ATCGCTATAATCCGGCGCCTACTGAGGGCCTATAGCTCAGTTGGTTAGAGCAGAGGACTCATAATCCTTTGGTCC
ACGGTTCAAGTCCGTGTGGGCCCACCAAACAAGAAAGCCGCGCAATGCGCGGCTTTCGCTTTTCTGAATCGTTCA
AGAAATAGCCCCTCCTTTCCTGTTCCGCCCATACAAGCTCAGTTAACGCTGGACAGGGCCGGGATGGCCCCTAGA
AGCCTCTTCTGCCCTCATCGCCGACCACCGATACAGACAGTCAACTTCTACTGCCATAGCGCAACTTCA

*ΔbluF*

CCATGAAGGCCTCGATCACCGATTCCGCCGCCGTGGCGGTTTCACACAGCACCACGCCGATGCCCTGGGTGCCTT
CCAGCACCTTGATCACCAACGGCGCGCCGTTGACCATCTCGATCAGGTCGGGAATGTCGTCCGGGGAATGGGCGA
AGCCGGTCACCGGCAGGCCGATGCCACGCCGCGACAGCAACTGCAGCGAGCGCAGCTTGTCCCGGGAGCGGGCGA
TGGCCACCGATTCGTTGAGGGGGAACACCCCCATCATTTCGAACTGGCGCAGCACCGCGCAGCCATAGAAGGTCA
CCGAGGCACCGATCCGCGGGATCACCGCATCGAAGCCTTCCAGGGGCTTGCCGCGGTAGTGGATCTGCGGCTTGT
GACTGGCGATGTTCATGTAGGCCCGCAGGGTATCGACCACCACCATTTCATGGCCACGCTCGGTGCCGGCCTCGA
CCAGACGACGAGTGGAATACAGACGCGGGTTACGCGACAGCACAGCGATCTTCATGCAACACCTGTGGCAGAGGT
AGTGGATACCGGGAACACCGGTTTGTCTTGTACGTACTTGATGCCCGGATTGACCACCAGTTGGCCGTCGATCAG
AGCCTTGGAACCCAGCAGCAGGCGATAGCGCATGGCCTTGCGGCAGGCGAGGGTGAACTCCACTCGCCACACCCG
ATCGCCCAGGGCCAGGGTGGTGCTGATCACGTAGCGCACCTGGGCCTGGCCGTTGGAGCTTTTAATGGTTTTCAT
CGCCACCAGCGGTGCTTCGCAGCGCCGGTGACGCAACTGCACCACACTGCCCAGGTGGGCATTGAAGCGCACCCA
CTGCTCACCGTCGCGCTCGAAGGGCTCGATCTCGGTGGCATGCAGGCTGGAGGTGCTGGCCCCGGTGTCGATCTT
GGCGCGCAGGCCGGCCACGCCCAGATCCGGGAGCGCCACCCACTCGCGCAGACCAACAACGGTCAAATGGTCAAA
AGTCTTCAATAAGGAAAATCCGCAATTAGGGGACTCCTCGCGTATCAGGGCGCAACCTGGGTTATTTTAGGTGAG
CGGCAGGGTTTATGAGAGCCCGCTTGCAGGCGGTAAAGTTGCGACATTTTTCAGCGAGGGGAATTCCAGTGGCGC
ACAAGCAGGAAGAAGAGGACAAGGTTCGTCTGGATAAATGGCTGTGGGCAGCGCGCTTCTACAAGACCCGGGCCC
TGGCCAAGACAGCGATTGAAAGCGGCAAGGTGCATCACCGGGGCGAGCGCTGCAAACCGGGCAAGGAGCCGCGGG
TAGGCGATGAATACCAGATTCGCGCCGGCTTCGACGAACGCACCGTGGTGGTCCAGGCCCTGTCCATCGTCCGCC
GTGGCGCTCCTGAAGCGCAGGCTCTGTACGCGGAAACCGAGGCCAGCATCGCCAAGCGCGAAGCCGCTGCGGCCC
AGCGCAAGGCTGGTGCCCTAGGAGTGAGCACCGACGGTAAACCGAGCAAGAAACAGCGCCGGGACCTGTTCAAGT
TCCACGGCAGCAACCACGAATGAAACCGTAGGAGCCGGCAAGCCGGCTCCTACGGGGCTGCTTAGCGCAGCAGGC

TGCTGGCGCGCACCACGCTCATGCGCGAGAGCACGGGCATCCGGCTCAACAGGTTGAACAGCGGGTGGGTCAGGC
GCAGCAGGAAGTTCGACATCCGCGCCGCGTACGGGGTGTAGTAGCCCCAGCCCAGGGCCAGCACCGCCAGCAGCA
CACCGCCGATGTAGTCGTCCTGGCCCCAGTGGGCGCCTGCCACCAGGCGCGGCATCATGAACAGCAGGGTCAGGC
CCCAGACGATCAGGAACTGGCCGATGCTGCGGCTGAAGATCCCCATGAACAGGCCCCAGATCAGCAGCACCGAAG
CGTGGTCGCCCGGGAAGCTCTGGCTGGAGCGGTCCTTGAGTTCCCAGGTCTTTTCCCAGCCCGGGAAGTAGTCGC
TCATGTGCACGGCGCCTTCCAGCACCATGGACGGGCTGTTGTGCTGCCAGTTC

### 4. Supplementary references

**de Lorenzo, V., Timmis, K.N. (1994)** Analysis and construction of stable phenotypes in gram-negative bacteria with Tn*5*- and Tn*10*-derived minitransposons. Methods Enzymol., Academic Press**:** 386-405.

**Dunn, R., McCoy, J., Simsek, M., Majumdar, A., Chang, S.H., Rajbhandary, U.L., Khorana, H.G. (1981)** The bacteriorhodopsin gene. *Proc. Natl. Acad. Sci. U. S. A.* **78**, 6744-6748. doi:10.1073/pnas.78.11.6744

**Gomelsky, M., Hoff, W.D. (2011)** Light helps bacteria make important lifestyle decisions. *Trends Microbiol.* **19**, 441-448. https://doi.org/10.1016/j.tim.2011.05.002

**Hmelo, L.R. *et al.* (2015)** Precision-engineering the *Pseudomonas aeruginosa* genome with two-step allelic exchange. *Nat. Protoc.* **10**, 1820-1841. 10.1038/nprot.2015.115

**Howell, C.R., Stipanovic, R.D. (1979)** Control of *Rhizoctonia solani* on cotton seedlings with *Pseudomonas fluorescens* and with an antibiotic produced by the bacterium. *Phytopathology* **69**, 480-482. 10.1094/Phyto-69-480

**Ihee, H., Rajagopal, S., Šrajer, V., Pahl, R., Anderson, S., Schmidt, M., Schotte, F., Anfinrud, P.A., Wulff, M., Moffat, K. (2005)** Visualizing reaction pathways in photoactive yellow protein from nanoseconds to seconds. *Proc. Natl. Acad. Sci. U. S. A.* **102**, 7145-7150. doi:10.1073/pnas.0409035102

**Kim, J.J., Sundin, G.W. (2001)** Construction and analysis of photolyase mutants of *Pseudomonas aeruginosa* and *Pseudomonas syringae*: contribution of photoreactivation, nucleotide excision repair, and mutagenic DNA repair to cell survival and mutability following exposure to UV-B radiation. *Appl. Environ. Microbiol.* **67**, 1405-1411. doi:10.1128/AEM.67.4.1405-1411.2001

**Losi, A., Gärtner, W. (2021)** A light life together: photosensing in the plant microbiota. *Photochem. Photobiol. Sci.* **20**, 451-473. 10.1007/s43630-021-00029-7

**Miller, W.G., Leveau, J.H.J., Lindow, S.E. (2000)** Improved *gfp* and *inaZ* broad-host-range promoter-probe vectors. *Mol. Plant-Microbe Interact.* **13**, 1243-1250. 10.1094/mpmi.2000.13.11.1243

**Okegbe, C., Fields, B.L., Cole, S.J., Beierschmitt, C., Morgan, C.J., Price-Whelan, A., Stewart, R.C., Lee, V.T., Dietrich, L.E.P. (2017)** Electron-shuttling antibiotics structure bacterial communities by modulating cellular levels of c-di-GMP. *Proc. Natl. Acad. Sci. U. S. A.* **114**, E5236-E5245. doi:10.1073/pnas.1700264114

**Quecine, M.C., Kidarsa, T.A., Goebel, N.C., Shaffer, B.T., Henkels, M.D., Zabriskie, T.M., Loper, J.E. (2016)** An interspecies signaling system mediated by fusaric acid has parallel effects on antifungal metabolite production by *Pseudomonas protegens* strain Pf-5 and antibiosis of *Fusarium* spp. *Appl. Environ. Microbiol.* **82**, 1372-1382. doi:10.1128/AEM.02574-15

**Silby, M.W. *et al.* (2009)** Genomic and genetic analyses of diversity and plant interactions of *Pseudomonas fluorescens*. *Genome Biol.* **10**, R51. 10.1186/gb-2009-10-5-r51

**Winkler, A., Udvarhelyi, A., Hartmann, E., Reinstein, J., Menzel, A., Shoeman, R.L., Schlichting, I. (2014)** Characterization of elements involved in allosteric light regulation of phosphodiesterase activity by comparison of different functional BlrP1 states. *J. Mol. Biol.* **426**, 853-868. https://doi.org/10.1016/j.jmb.2013.11.018

**Wu, L., McGrane, R.S., Beattie, G.A. (2013)** Light regulation of swarming motility in *Pseudomonas syringae* integrates signaling pathways mediated by a bacteriophytochrome and a LOV protein. *mBio* **4**, 10.1128/mbio.00334-00313. doi:10.1128/mbio.00334-13

**Yan, Q., Philmus, B., Chang, J.H., Loper, J.E. (2017)** Novel mechanism of metabolic co-regulation coordinates the biosynthesis of secondary metabolites in *Pseudomonas protegens*. *eLife* **6**, e22835. 10.7554/eLife.22835

**Yan, Q., Philmus, B., Hesse, C., Kohen, M., Chang, J.H., Loper, J.E. (2016)** The rare codon AGA is involved in regulation of pyoluteorin biosynthesis in *Pseudomonas protegens* Pf-5. *Front. Microbiol.* **7**, 497. 10.3389/fmicb.2016.00497
